# KLRG1 identifies circulating cytotoxic CD4 T cells with selective anti-tumor function in human cancer

**DOI:** 10.64898/2026.04.13.718113

**Authors:** Mara Cenerenti, Josep Garnica, Margaux Saillard, Paul Gueguen, Benita Wolf, Florent Lemaitre, Romina Marone, Yen-Cheng Liu, Anthony Cornu, Alexandre Dumez, Maria Giulia Sardiello, Robert Pick, Stéphane Jemelin, Laurence de Leval, Sabina Müeller, Salvatore Valitutti, Christoph Scheiermann, Ronjon Chakraverty, Hatice Altug, Daniel E. Speiser, Jean Villard, Lukas T. Jeker, Virginie Hamel, Pedro Romero, Santiago Carmona, Camilla Jandus

## Abstract

Given the critical role of CD4 T cells in anti-tumor immunity, strategies to harness these cells for cancer immunotherapy are gaining increasing interest. Historically overshadowed by CD8 T cells, cytotoxic CD4 T cells can directly kill MHC class II-expressing tumor cells. However, the defining molecular signature and the mechanisms underlying their cytolytic activity remain poorly understood, particularly in cancer patients.

Here, using *ex vivo* single-cell transcriptomic and spatial analyses of CD4 T cells from paired blood and tumor samples of melanoma patients, we identified Killer Cell Lectin-Like Receptor G1 (KLRG1) as a defining surface marker of cytotoxic CD4 T cells. The CD4^+^ KLRG1^+^ T cell subset was notably enriched among circulating cells compared with tumor-infiltrating populations, which were instead enriched in T follicular helper (Tfh) states.

Functionally, KLRG1^+^ CD4 T cells expressed elevated levels of cytotoxic genes and exhibited superior tumor-killing capacity compared with their KLRG1^-^ counterparts. We demonstrated that their cytotoxicity is granulysin-dependent, as confirmed by CRISPR/Cas9-mediated gene deletion. Mechanistically, CD4 T cells spared MHC class II^+^ cells lacking the KLRG1 ligands CD324 and CD325, such as professional antigen-presenting cells (APCs), indicating that cytotoxicity was selectively directed towards tumor cells while preserving immune cells.

Finally, by investigating how the tumor microenvironment may impair CD4 T cell cytotoxicity, we showed that tumor-derived factors, including interleukin-6 (IL-6), are key drivers promoting the transition of cytotoxic CD4 T cells toward a Tfh phenotype.

In summary, our findings define KLRG1 as a defining cell surface marker of cytotoxic CD4 T cells in cancer patients, as well as a key regulator that protects MHC class II^+^ APCs. Moreover, targeting the IL-6 signalling pathway may enhance CD4 T cell anti-tumor cytotoxicity, offering new avenues for cancer immunotherapy.

## INTRODUCTION

T cells are key coordinators of the anti-cancer immunity, and T cell targeted immunotherapies have shown remarkable efficacy in patients [1–5]. However, restoring CD8 T cell function remains challenging due to persistent cellular dysfunction or exhaustion [6, 7] and their efficacy is further compromised by tumor MHC class I loss, a common mechanism of resistance [8, 9]. These limitations highlight the need for CD8/MHC I-independent therapeutic strategies.

CD4 T cells have gained attention not only for their helper functions in sustaining antigen presenting cells (APCs) through activation signals, but also for their capacity to directly eliminate tumor cells presenting tumor-associated or neoantigens in the context of MHC class II [10, 11]. Single-cell RNA (scRNA-seq) studies have identified distinct populations of CD4 T cells expressing cytotoxic programs across multiple human cancer types [12–25].

Moreover, cytotoxic CD4 T cells (CD4 CTLs) are induced in patients with advanced melanoma following immune checkpoint blockade [18] and adoptive cell therapy (ACT) [26–28], as well as in leukaemia patients treated by CD19 chimeric antigen receptor (CAR) T cells, as assessed ten years after treatment [29]. Preclinical studies have demonstrated that CD4 CTLs can directly eliminate and eradicate tumors in an MHC class II-restricted manner [30]. Consistent with these findings, tumor-reactive cytotoxic CD4 T cells have been shown to expand *in vivo* and eradicate established melanoma following the transfer of naïve CD4 T cells into lymphopenic hosts [31]. Furthermore, in murine melanoma models, the transfer of a small number of CD4 T cells into lymphopenic mice, combined with CTLA-4 blockade [30] or CD137 agonist immunotherapy [32], leads to potent tumor rejection independent of other immune cell populations and in an MHCII-restricted manner. Finally, we recently demonstrated the presence of highly cytotoxic, tumor-reactive CD4 T cells both in melanoma patients, characterised by conserved and preferential TCR pairing, supporting their broad applicability in TCR-engineered ACT across different cancer types, both adult and pediatric [33].

However, whether CD4 CTLs consistently promote anti-tumor responses or may, under certain conditions, contribute to tumor progression remains unclear. Their functional plasticity likely depends on tumor type, disease stage, and the immunological context of the tumor microenvironment.

With regard to the cytotoxic mechanisms potentially employed by CD4 T cells, several effector molecules have been proposed, including GZMB, GZMA, and PRF1, in the context of murine and human viral infections, aging, and cancers [12–14, 34, 35]. Phenotypically, CD4 CTLs are described as effector or effector-memory, antigen-experienced cells that have downregulated co-stimulatory receptors such as CD27 and CD28 [36, 37].

Recent work has provided novel insight into the developmental origins and cytolytic potential of CD4 T cells. One study identified a stem-like subset of CD4 T cells (TSCM-CTLs) poised for cytotoxic differentiation, with transcriptional and functional profiles similar to CD8 CTLs, highlighting a defined pathway for CD4 cytotoxicity in humans [38]. These findings support the concept that CD4 CTLs are both functionally potent and developmentally programmed for cytotoxic responses in specific immunological contexts.

Despite these advances, a clear cell surface marker that reliably identifies tumor-specific cytotoxic CD4 T cells directly *ex vivo* in cancer patients is still lacking, and the precise cytotoxic mechanisms employed by these cells in patients remain incompletely defined.

Here, we identify KLRG1 as a defining marker of cytotoxic CD4 T cells that utilise a granulysin- dependent killing mechanism. We demonstrate that these KLRG1⁺ CD4 T cells selectively target tumor cells while sparing APCs, which lack cadherins, the ligands for KLRG1, and are therefore protected from killing. This selective cytotoxicity provides a mechanistic basis for the immune privilege of APCs, a concept long proposed to be essential for maintaining immune integrity, and further supports the therapeutic potential of harnessing CD4 T cells as cytotoxic effectors.

Furthermore, we show that the cytolytic activity of these cells is attenuated within the tumor microenvironment, likely due to IL-6–mediated signalling, and that interference with this pathway may restore their anti-tumor function.

Together, our findings provide new insights into the identification, regulation, and therapeutic potential of cytotoxic CD4 T cells in cancer immunotherapy.

## MATERIALS AND METHODS

### Lymphocyte isolation from peripheral blood and tumor tissues

Peripheral blood mononuclear cells (PBMC) and tumor-infiltrated lymph nodes (TILN) from patients with stage III/IV melanoma (Supp Fig.1A) were obtained from the Department of Oncology, University Hospital (CHUV), Lausanne, Switzerland (NCT00112229, NCT00002669 and NCT00002763) under the approval of the Lausanne University Hospital’s Institutional Review Board. Blood was diluted with Dulbecco’s Phosphate Buffered Saline 1X (DPBS, Sigma-Aldrich) and PBMC were purified by density gradient centrifugation using Lymphoprep^TM^ (StemCell). TILN were obtained after surgery from melanoma patients from the Department of Oncology, University Hospital (CHUV), Lausanne, Switzerland (EC 2023-00186). Resected tumors were enzymatically digested and were used as starting material for TILN generation, as previously reported [39].

### Generation of antigen-specific CD4 and CD8 T cell clones

PBMC and TILN cells (in short TILN) from HLA-DRB1*07:01 melanoma patients were thawed and CD4 T cells were purified by positive selection using MACS isolation microbeads (Miltenyi). These cells were then stimulated with tumor associated antigen NY-ESO-1_87-99_ (2 µM) and with autologous irradiated (30 Gray) CD4 negative cells in Roswell Park Memorial Institute medium (RPMI) containing 8% of human serum (AB^+^, male; purchased from BSZ Basel, Switzerland), 1% L-Glutamine (Gibco), 1% (vol/vol) nonessential amino acids (Gibco), 1% Na-Pyruvate (Gibco), 0.1% 2β-mercaptoethanol (Gibco), 1% Kanamycin (Gibco), 10mM Hepes Buffer (Gibco), 1% penicillin (Gibco) and 1% streptomycin (Gibco) (named as “RPMI 8% HS”). After 2 days of culture, 100 µL of RPMI 8% HS media containing a final concentration of 100 IU/mL of human recombinant interleukin-2 (IL-2) (Proleukine) was added. After 10 days of *in vitro* expansion, NY-ESO-1_87-99_-specific CD4 T cells were stained with fluorescent peptide-MHC class II (pMHCII) multimers (produced in house) with an optimized protocol developed in house [40]. Cells were first resuspended in RPMI 8% HS with 5 mM N-Acetyllactosamine (LacNac, Sigma-Aldrich) for 2 hours at 37°C and then resuspended in DPBS 1X containing 50 nM Dasatinib (Axon) for 30 min at 37°C. Multimer staining was performed using a PE-pMHCII/HLA-DRB1*07:01/NY-ESO-1_87-99_ multimer for 25 min at 37°C and the surface staining with an APC anti-CD3 (clone UCHT1, 1:50 dilution, Biolegend) and PerCP-Cy5.5 anti-CD4 antibodies (clone OKT4, 1:200 dilution, Biolegend) and LIVE/DEAD Fixable Near-IR Dead Cell Stain (Thermo Fisher Scientific) for 20 min at RT. Finally, a mouse anti-PE antibody (clone PE001, 1:50 dilution, Biolegend) was added for 20 min at 4°C. Positive cells were sorted using a FACSAria Fusion (BD). Individual, sorted, NY-ESO-1_87-99_ CD4 T cells were stimulated with 1 μg/mL phytohemagglutinin (PHA, Thermo Fisher Scientific) in the presence of irradiated (30 Gray) allogeneic PBMC feeder cells and IL-2 (100 IU/mL) and plated in Terasaki plates (Greiner Bio-One) for clone generation.

CD8 T cell clones were generated with the same protocol using PBMC from HLA-A*02:01 melanoma patients stimulated with the tumor associated antigen Melan-A_26-35(A27L)_ peptide and sorted with a PE- pMHCI/HLA-A*02:01/MelanA_26-35(A27L)_ multimer.

### Synthetic peptides

Peptides were synthesized by the Peptide and Tetramer Core Facility, UNIL-CHUV, Epalinges, Switzerland using INTAVIS synthesizers. All peptides were >90% pure as indicated by analytic mass spectrometry and high-performance liquid chromatography. The lyophilized peptides were diluted in pure dimethylsulfoxide (DMSO) at 10 mM and stored at −80°C.

### Tumor cell lines

Melanoma cell lines were established from metastatic melanoma patients treated at the Department of Oncology, University Hospital (CHUV), Lausanne, Switzerland (Me215, Me275, Me252, T333A, T672E, T311B, T618A), and by the laboratory of Prof. Maria Pia Protti, San Raffaele Scientific Institute, Milano, Italy (GEFI). Tumor cell lines were cultured in RPMI containing 10% of Fetal Calf Serum (FCS, Gibco), 10 mM Hepes Buffer (Gibco), 0.55 mM L-Arginine (Sigma-Aldrich), 0.24 mM L-Asparagine (Sigma-Aldrich), 1.5 mM Glutamine (Gibco), 0.05 mg Ciproxin (Bayer), 1% penicillin (Gibco) and 1% streptomycin (Gibco) (named as “RPMI 10% FCS”).

### Generation of CIITA-transduced tumor cell lines

6-hour before transfection, 293T cells were seeded at 1.25 x 10^6^ in 2 mL RPMI 10% FCS media/well in a 6-well plate. 293T cells were transfected with 2.5 µg total DNA (divided as 0.282 µg pVSVG, 0.846 µg of R874, and 1.125 µg of plasmid containing the human CIITA isoform 3 gene), using a mix of Lipofectamine 2000 (Invitrogen) and OptiMEM media (Gibco). The viral supernatant was harvested 48 hours post-transfection and the supernatant was used directly on melanoma cell lines. MHC class II positive cells were sorted using a FACSAria Fusion (BD) after staining with a FITC anti-HLA-DR/-DQ/- DP antibody (clone Bu26, 1:10 dilution, Abcam).

### Contact frequency and duration of individual tumor cell-T cell in picowell arrays

Data generated in [14] were analysed using an artificial intelligence-assisted approach with a customized version of Deeplabcut, an open-source toolkit for marker less pose estimation based on transfer learning with deep neural networks. In this study, we adapted this toolkit for cell segmentation to identify cell boundaries in the provided time-lapse fluorescence images. The deep neural network was trained using cropped frames, each containing 3x3 picowells with various compositions of effector-target cells, with features such as picowell boundaries, alive tumor cells, lysed tumor cells, alive T cells, and lysed T cells labelled accordingly. Subsequently, the time-lapse videos were analysed by the trained network to predict the pixel coordinates of all detections in each frame. The predicted coordinates were then processed in Python to analyse cell contact events, defined as sets of consecutive frames where the distance between the closest pixels of two cell detections was within a threshold distance of 10 pixels. The contact frequency was determined as the number of contact events during the measurement time course, while the duration of contact was calculated as the time duration of one contact event (number of consecutive frames × time interval between frames). In this study, only the cases involving one effector cell and one target cell in each well were analysed.

### Cryo-fixation coupled to Ultrastructure Expansion Microscopy

The cell seeding, co-culture and cryo-fixation [41, 42] and ultrastructure expansion microscopy (UExM) [43, 44] was performed as previously described for T cell tumor cell pairs.

In brief, melanoma (GEFI-CIITA) cells were seeded on 12 mm poly-D-lysine–coated coverslips (0.1 mg/ml, overnight, 4°C) at 3 x 10⁵ cells per coverslip. After 30 min of adhesion, cells were pulsed for 1 hour with peptide (MelanA_26-35(A27L)_: 2 μM for CD8 T cells; NY-ESO-1_87-99_: 3 μM for CD4 T cells). 3 x 10⁵ T cells were added after washing. Fixation times were guided by prior time-lapse imaging: CD8 T cells were fixed after 30 min, and CD4 T cells after 3 hours of co-culture.

For cryo-fixation, coverslips were blotted and plunge-frozen in liquid ethane (−170°C) using fine tweezers. Samples were transferred into acetone (pre-chilled with liquid nitrogen) containing 0.1% paraformaldehyde (PFA, Sigma-Aldrich) and 0.02% glutaraldehyde (GA, Sigma-Aldrich), and incubated overnight on dry ice under gentle agitation. After gradual warming to ∼0 °C, samples were rehydrated through ethanol–water steps and stored in PBS. Cryo-fixed cells were pre-incubated for 3 hours at 37°C in 2% acrylamide and 1.4% formaldehyde. Gelation followed in a monomer solution (19% sodium acrylate, 10% acrylamide, 0.1% bis-acrylamide, with 0.5% tetramethylethylenediamine (TEMED) and ammonium persulfate (APS), with polymerization at 37°C for 1 hour after a 5 min ice pre-incubation. Gels were denatured at 95°C for 1.5 hours, washed in ddH₂O, and the expansion factor was determined by comparing gel diameter to the original coverslip size. Expanded gels were incubated in PBS, stained with primary antibodies in 2% BSA–PBS at 37°C for 3 hours, then washed and stained with secondary antibodies. After final PBS–Tween washes, gels were expanded in ddH₂O. If required, NHS-ester staining was performed before the final expansion. The following primary antibodies and dyes were used in this study: anti-α-tubulin (clone scFv-F2C, 1:250 dilution, ABCD antibodies), anti-β-tubulin (clone scFv-S11B, 1:250 dilution, ABCD antibodies), anti-β-actin (clone 7D2C10, 1:250 dilution, Proteintech), anti-α-tubulin (polyclonal, 1:250 dilution, Abcam), anti-LAMP1 (polyclonal, 1:100 dilution, R&D Systems), anti-GZMB (clone 351927, 1:200 dilution, R&D Systems), anti-PRF1 (polyclonal, 1:250 dilution, Proteintech), anti-CD8 (clone 1G2B10, 1:500 dilution, Proteintech), anti-CD4 (clone EPR6855, 1:250 dilution, Abcam), anti-GNLY (clone E2T3D, 1:250 dilution, Cell Signaling Technology), anti-NHS-Ester–Alexa Fluor 405 (40 ng/ml, Thermo Fisher Scientific). The following secondary antibodies were used: anti-mouse Alexa Fluor 568 (1:400 dilution, Invitrogen), anti-mouse Alexa Fluor 488 (1:400 dilution, Invitrogen), anti-rabbit Alexa Fluor 488 (1:400 dilution, Invitrogen), anti-rabbit Alexa Fluor 568 (1:400 dilution, Invitrogen), anti-sheep Alexa Fluor 488 (1:400 dilution, Invitrogen), anti-rabbit Alexa Fluor Cy3 (1:250 dilution, Jackson ImmunoResearch), anti-mouse Alexa Fluor 647 (1:250 dilution, Jackson ImmunoResearch), anti-rabbit Alexa Fluor 647 (1:400 dilution, Thermo Fisher Scientific).

### Image acquisition

Gel fragments were mounted on poly-D-lysine–coated 24 mm precision coverslips (No. 1.5H, Marienfeld) for imaging, as described in [42]. Images were acquired using a Leica DMi8 Thunder widefield microscope (63X, 1.4 NA oil) controlled via LAS X software (Leica Microsystems). Widefield images were processed using large volume computational clearing (LVCC) with adaptive settings and water as mounting medium. Typical voxel sizes were 0.1013 × 0.1013 × 0.2131 µm for widefield.

### 3D granule segmentation

Lytic granules containing PRF1 and GZMB were segmented in 3D using LABKIT in Fiji, as described in [42]. Pixel classifiers were trained on annotated 3D image stacks from three independent experiments, using a set of 3D features (e.g., Gaussian blur, Laplacian of Gaussian, structure tensor). Classifiers were refined as needed and applied via a custom Fiji macro to segment granules in primary T cells.

Synaptic landmarks were identified using orthogonal views, and total PRF1 and GZMB intensities were quantified in maximum intensity projections. Segmented probability maps were thresholded (Otsu method) to create binary masks, filtered for granule size (>12 voxels), and analysed with the 3D Object Manager for intensity and morphology.

Granule colocalization was assessed using 3D MultiColoc in Fiji, and polarization indices were calculated based on granule distance from the synaptic dome. Final data analysis was performed in R, and graphs were generated using GraphPad Prism (version 10).

### Single-cell (sc) RNA sequencing (RNAseq) analysis

PBMC and TILN were thawed, washed and stained. Multimer staining was performed using a PE- pMHCII/HLA-DRB3*02:02/NY-ESO-1_123-137_, BV421-pMHCII/HLA-DRB1*07:01/NY-ESO-1_87-99_, PE-pMHCII/HLA-DPB1*04:01/MAGE-A3_243-258_, for 20 min at RT and the surface staining with BUV395 anti- CD3 (clone UCHT1, 1:400 dilution, BD), PerCP-Cy5.5 anti-CD4 (clone OKT4, 1:200 dilution, Biolegend) antibodies and LIVE/DEAD Zombie Green Fixable Viability Kit (1:1000, Biolegend) for 20 min at RT. Cells were washed and 1 mg of hashtags (TotalSeq^TM^-C anti-human antibody, Biolegend) were added for 30 min at 4°C. Cells were isolated using FACSAria Fusion (BD). Sorted cells were loaded onto the 10x Genomics Chromium Controller. cDNA amplification and gene expression library preparation were performed using the Chromium Next GEM Single Cell 5’ Kit v2 and the 5’ Feature Barcode Kit. The Library Construction Kit was used for final library construction. All procedures followed the manufacturer’s protocols (10x Genomics). Library quantification and quality control were carried out using a Qubit Fluorometer (Thermo Fisher Scientific) and a TapeStation system with DNA High Sensitivity chips (Agilent Technologies). Libraries were sequenced on an Illumina NovaSeq 6000 using a paired-end 28x90 read configuration.

### Ex vivo staining of KLRG1^+^ cells

For multimer staining, PBMC were stained directly *ex vivo* using BV421-pMHCII/HLA-DRB1*07:01/NY- ESO-1_87-99_ multimer, PE/Dazzle594 anti-gp130 (clone 2E1B02, 1:800 dilution, Biolegend), BV605 anti- KLRG1 (clone 13F12F2, 1:100 dilution, eBioscience), BUV395 anti-CD3 (clone UCHT1, 1:400 dilution, BD), BUV737 anti-CD4 (clone SK3, 1:800 dilution, BD) antibodies and LIVE/DEAD Fixable Near-IR Dead Cell Stain kit (Thermo Fisher Scientific) at RT for 20 min. All samples were acquired at LSR Fortessa II (BD) and analysed using FlowJo software (v.10.10.0).

For intracellular staining, PBMC were stained *ex vivo* using ECD anti-CD45RA (clone 2H4LDH11LDB9, 0.3:50 dilution, Beckman Coulter), PE-Cy7 anti-CD25 (clone BC96, 0.1:50 dilution, Biolegend), BV421 anti-CCR7 (clone A019D5, 1:50 dilution, Biolegend), BV605 anti-KLRG1 (clone 13F12F2, 1:100 dilution, eBioscience), BV785 anti-CD127 (clone A019D5, 1:50 dilution, Biolegend), BUV395 anti-CD3 (clone UCHT1, 1:400 dilution, BD), BUV737 anti-CD4 (clone SK3, 1:800 dilution, BD) antibodies and LIVE/DEAD Fixable Near-IR Dead Cell Stain kit (Thermo Fisher Scientific) at RT for 20 min. Then cells were fixed and permeabilized for 30 min at RT with Foxp3/Transcription Factor Staining Buffer Set (Invitrogen) and stained with FITC anti-PRF1 (clone dG9, 1:50 dilution, Biolegend), PerCP-Cy5.5 anti-GZMK (clone G3H69, 1:100 dilution, eBioscience), PE anti-GNLY (clone DH2, 1:50 dilution, eBioscience), APC anti-GZMM (clone 50-9774-42, 1:50 dilution, eBioscience), AlexaFluor700 anti-GZMA (clone CB9, 0.1:50 dilution, Biolegend), BV510 anti-GZMB (clone GB11, 1:100 dilution, BD) antibodies for 30 min at RT. All samples were acquired at LSR Fortessa II (BD) and analysed using FlowJo software (v.10.10.0).

### Ex vivo sorting of KLRG1^+^ cells

PBMC were stained *ex vivo* using Alexa Fluor 700 anti-CD45RA (clone HI100, 1:100 dilution, BD), FITC anti-CD3 (clone UCHT1, 1:200 dilution, Biolegend), APC anti-CD4 (clone A161A1, 1:400 dilution, Biolegend), BV421 anti-CCR7 (clone G043H7, 2:50 dilution, Biolegend), BV785 anti-CD127 (clone A019D5, 1:50 dilution, Biolegend), BV510 anti-CD25 (clone BC96, 1:100 dilution, Biolegend), PE anti- KLRG1 (clone 13F12F2, 1:400 dilution, eBioscience) and LIVE/DEAD Fixable Near-IR Dead Cell Stain kit (Thermo Fisher Scientific) at RT for 20 min, then sorted using FACSAria Fusion (BD).

### RNA purification and RT-qPCR

Total RNA was isolated using the TRIzol reagent according to the manufacturer’s instructions (Invitrogen). The final preparation of RNA was considered DNA and protein free if the ratio of spectrophotometer (NanoDrop 2000, Thermo Fisher Scientific) readings at 260/280 nm was ≥1.7. Isolated mRNA was reverse-transcribed using the iScript^TM^ cDNA Synthesis Kit (Bio-Rad) according to the manufacturer’s protocol. The real-time quantitative polymerase chain reaction (RT-qPCR) was carried out in the QuantStudio^TM^ 6 Pro Real-Time PCR System (Applied Biosystems) with specific primers (Microsynth AG Switzerland) (hCST7, 5’-TTGTTCAAGGAGTCCCGCATC-3’ and 5’-GTCACAGTCATCCAGACGCA-3’; hGNLY, 5’-CCTGTCTGACGATAGTCCAAAAA-3’ and 5’-GACCTCCCCGTCCTACACA-3’; hGZMA, 5’-CCCTATCCATGCTATGACCCA-3’ and 5’-AGTATCGGACCAAGATGCACTAT-3’; hGZMB, 5’-TACCATTGAGTTGTGCGTGGG-3’ and 5’-GCCATTGTTTCGTCCATAGGAGA-3’; hGZMH, 5’-CTGGCTGGGGTTATGTCTCAA-3’ and 5’-GGCTACGTCCTTACACACGAG-3’; hGZMK, 5’-GGGGCTTATATGACTCATGTGTG-3’ and 5’-GTGGATCAATCAGAACACCTCC-3’; hNKG7, 5’-GATTGCTTTGAGCACCGATTTC-3’ and 5’-GTAGCCTGATATGATGTCCCCA-3’; hPRF1, 5’-GACTGCCTGACTGTCGAGG-3’ and 5’-TCCCGGTAGGTTTGGTGGAA-3’; hBCL6, 5’-ACACATCTCGGCTCAATTTGC-3’ and 5’-AGTGTCCACAACATGCTCCAT-3’; hCXCR5, 5’-CACGTTGCACCTTCTCCCAA-3’ and 5’-GGAATCCCGCCACATGGTAG-3’; hICOS, 5’-ACAACTTGGACCATTCTCATGC-3’ and 5’-TGCACATCCTATGGGTAACCA-3’; hPD1, 5’-CCAGGATGGTTCTTAGACTCCC-3’ and 5’-TTTAGCACGAAGCTCTCCGAT-3’; hIL11RA, 5’-CTGGGCTAGGGCATGAACTG-3’ and 5’-CTGGGACTCCAAGTGCAAGA-3’; hIL6-R, 5’-CCCCTCAGCAATGTTGTTTGT-3’ and 5’-CTCCGGGACTGCTAACTGG-3’; hLIFR, 5’-TGGAACGACAGGGGTTCAGT-3’ and 5’-GAGTTGTGTTGTGGGTCACTAA-3’) using KAPA SYBR FAST Kit (Kapa Biosystems). All RT-qPCR reactions were performed in duplicate and amplified simultaneously with a nontemplate control blank for each primer pair to control for contamination or for primer dimerization, as well as a housekeeping gene (RPS18) to normalize and determine the Ct values, using the 2^−DCt^ formula.

### Redirected killing assay and LDH cytotoxicity assay

In the redirected killing assay, the FcγR^+^ murine mastocytoma cell line P815 (ATCC) was used as target cell line. P815 cells were incubated with CD4 T cells at 10:1 (E:T) ratio for 4 hours at 37°C in the presence of anti-CD3 antibody (3.2 mg/ml, clone OKT3, Biolegend). At the end of the incubation, supernatants were collected and mixed with Reaction mixture (CyQUANT™ LDH Cytotoxicity Assay Kit, Thermo Fisher Scientific) for 30 min at RT in the dark. Stop solution was added and the absorbances at 490 nm and 680 nm were measured to determine LDH activity (Ledetect96). The percentage of specific lysis was calculated as follows: (experimental – effector spontaneous release – target spontaneous release) / (target maximal release – target spontaneous release) x 100.

For KLRG1 blocking experiment, KLRG1^+^ CD4 T cells were treated with anti-KLRG1 antibody (clone 14C2A07, Biolegend) at 10 µg/ml for 20 min in 100 µl of DPBS 1X and co-cultured with target cells for 4 hours, then LDH cytotoxicity assay was performed.

For IL6-R blocking experiment, anti-human IL6-R antibody (tocilizumab, MedChemExpress) was added at 50 µg/ml during the co-culture for 48 hours with the target cells, then LDH cytotoxicity assay was performed.

### Genetic engineering of human primary CD4 T cells by CRISPR/Cas9

Protocols for human CRISPR/Cas9-mediated genome engineering are based on [45]. *Streptococcus pyogenes* Cas9 fused to two SV40 nuclear localization signals was produced in *E. coli* and purified using a combination of metal-affinity and size-exclusion chromatography methods. In short, Cas9 RNPs were freshly generated prior to each electroporation. Thawed crRNA and tracrRNA (purchased from IDT Technologies; at 200 µM) were mixed in a 1:1 M ratio (120 pmol each), denatured at 95°C for 5 min, and annealed at RT for 10–20 min to complex an 80 µM gRNA solution. Polyglutamic acid (15–50 kDa at 100 mg/ml; Sigma-Aldrich) was added to the gRNA (crRNA+tracrRNA) in a 0.8:1 volume ratio [46]. To complex RNPs, 60 pmol recombinant Cas9 (Protein and Peptides Facility UNIGE; at 40 µM) was mixed with the gRNA (molar ratio Cas9 : gRNA = 1 : 2) and incubated for 20 min at RT. Electroporation was performed with the 4D-Nucleofector system (Lonza) with Program DN-100. T cells were resuspended in 20 µl Lonza-supplemented P3 buffer. HDRT (3–4 µg) and RNPs (60 pmol) were mixed separately and incubated for 5 min. The cells were added to the mix and the total volume was transferred to 16-well Nucleocuvette Strips. Immediately following electroporation, 80 µl of prewarmed RPMI 8% HS media was added to each cuvette and incubated at 37°C. After 20 min, the cells were transferred into 48-well culture plates and replenished with IL-2 (50 IU/ml), IL-7 (10 ng/ml) and IL-15 (10 ng/ml) (Peprotech). Control T cells were electroporated with an Alt-R CRISPR Negative Control crRNA #1 (IDT Technologies). PCRs were performed and sent for Sanger sequencing (Microsynth AG Switzerland). KO efficiency was quantified using TIDE (available at http://tide.nki.nl).

### Cadherin expression quantification

PBMC, TILN and tumor cells were stained with APC-Fire 700 anti-CD324 (clone 67A4, 1:400 dilution, Biolegend) and PE-Cy7 anti-CD325 (clone 8C11, 1:25 dilution, Biolegend) antibodies at RT for 20 min. All samples were acquired at LSR Fortessa II (BD) and analysed using FlowJo software (v.10.10.0).

### Specific lysis quantification by 7AAD staining

PBMC from melanoma patients were incubated with an HLA-matched antigen-specific CD4 CTL clone, previously stained with CellTrace Far Red (1:1000 dilution, Invitrogen), at 1:1 (E:T) ratio for 24 hours at 37°C in the presence of the specific peptide (3 µM). To analyse B cells, monocytes and dendritic cells, cells were stained with FITC anti-CD3 (clone UCHT1, 1:200 dilution, Biolegend), PerCPCy5.5 anti- CD19 (clone HIB19, 1:50 dilution, Biolegend), PE anti-CD16 (clone 3G8, 0.5:50 dilution, Beckman Coulter), PE-CF594 anti-CD14 (clone MφP9, 1:200 dilution, BD), Alexa700 anti-CD11c (clone B-ly6, 0.3:50, BD), 7AAD (Invitrogen), BV510 anti-CD1c (clone L161, 2:50 dilution, Biolegend), BV605 anti- CD141 (clone M80, 1:200 dilution, Biolegend), BV650 anti-CD123 (clone 6H6, 1:100 dilution, Biolegend), BV711 anti-HLA-DR (clone L243, 1:200 dilution, Biolegend), BV785 anti-CD303 (clone 201A, 1:25 dilution, Biolegend) antibodies. All samples were acquired at LSR Fortessa II (BD) and analysed using FlowJo software (v.10.10.0).

### Multiplex fluorescent IHC staining of melanoma tissue sections

Immunostainings were performed on 5 µm formalin-fixed paraffin-embedded tissue sections from 5 melanoma patients, provided by the Institute of Pathology at the Lausanne University Hospital, CHUV. Following deparaffinization and heat-induced antigen retrieval, tissue sections from 5 melanoma patients (LAU706, LAU380, LAU1417, LAU372, LAU1397) were incubated in blocking buffer (PBS supplemented with 20% goat serum and 0.5% Triton X-100) for 2 hours at room temperature. Sections were then incubated overnight at 4°C with primary antibodies diluted 1:100 in PBS containing 2% goat serum to label KLRG1⁺/CD4⁺ T cells and CD325⁺/SOX10⁺ tumor cells. Primary antibodies included mouse anti-human SOX10 (clone BC34, Biocare Medical), goat anti-human CD4 Alexa Fluor 488 (polyclonal, R&D Systems), sheep anti-human CD325 (polyclonal, R&D Systems) and rabbit anti-human KLRG1 (clone 2388C, R&D Systems). After washing, sections were incubated for 2 hours at room temperature with the corresponding secondary antibodies diluted 1:1000: goat anti-mouse Alexa Fluor 405 (Thermo Fisher Scientific), donkey anti-sheep Alexa Fluor 568 (Thermo Fisher Scientific) and goat anti-rabbit Alexa Fluor 647 (Thermo Fisher Scientific). Imaging was performed using an upright spinning disk confocal microscope (Axio Examiner Z1 Advanced Microscope Base, Zeiss) equipped with a CSU-X1 confocal scanner unit (Yokogawa Electric Corporation). Fluorochromes were excited using 405, 488, 561 and 640 nm laser lines (LaserStack v4 Base, 3i). Fluorescence emission was collected with a 20×/1.0 NA water-immersion objective (W Plan Apochromat, Zeiss), appropriate band- pass emission filters (Semrock) and an electron-multiplying CCD camera (Evolve 512, 10 MHz back-illuminated, Photometrics). Three-dimensional image stacks were acquired by sequential imaging of 25 fields of view per sample using a motorized XY stage (ProScan, Prior). Image acquisition, deconvolution using the “No neighbours” algorithm and generation of maximum-intensity projections were performed using SlideBook software (v6.0.17, 3i). Montage images from contiguous fields were generated using the Fiji Grid/Collection stitching plugin [47]. Uneven background illumination was corrected using a rolling-ball subtraction algorithm implemented in the Fiji Subtract Background plugin. Identification and segmentation of KLRG1⁺/CD4⁺ T cells and CD325⁺/SOX10⁺ tumor cells were performed using morphological reconstruction implemented in the ImageJ plugin MorphoLibJ [48].

### GeoMx DSP RNA profiling in situ hybridization

GeoMx DSP combines standard immunofluorescence techniques with digital optical barcoding technology to perform highly multiplexed, spatially resolved profiling experiments [49]. DNA oligonucleotide probes were designed to bind mRNA targets. From 5′ to 3′, they each comprised of a 35- to 50-nucleotide target complementary sequence, an ultraviolet (UV) photocleavable linker and a 66-nucleotide indexing oligonucleotide sequence containing a unique molecular identifier (UMI), RNA ID sequence and primer binding sites. FFPE tissue section of 5 µm from 5 melanoma patients (i.e., LAU706, LAU380, LAU1417, LAU372, LAU1397) were mounted on positively charged histology slides. Sections were baked at 65°C for 1 hour. Slides were deparaffinized in 3 xylol baths of 5 min, then rehydrated in ethanol gradient from 100% EtOH, 2 baths of 5 min followed by 95% EtOH, 5 min. Slides were then washed in PBS 1X. Antigen retrieval was done in Tris-EDTA pH 9.0 buffer at 100°C for 20 min at low pressure. Slides were first deep into hot water 10 sec to be then deep into Tris-EDTA buffer. Cooker vent stay open during the procedure to ensure low pressure and reach 100°C. Slides were then wash into PBS 1X, and incubated into proteinase K in PBS (1 mg/ml) for 15 min at 37°C and washed again in PBS 1X. Tissue are post-fixed in 10% neutral-buffered formalin 5 min, washed 2 times 5 min in NBF stop buffer (0.1M Tris Base, 0.1M Glycine) and finally one time in PBS 1X. The mix of Whole Transcriptome Atlas probes (WTA, Nanostring) was dropped on each section and covered with HybriSlip Hybridization Covers. Slides were then put for hybridization overnight at 37°C in a Hyb EZ II hybridization oven (Advanced cell Diagnostics). The day after, HybriSlip covers were gently removed and 25 min stringent washes were performed twice in 50% formamide and 2X saline-sodium citrate (SSC) at 37 °C. Tissues were washed for 5 min in 2X SSC, then blocked in Buffer W (Nanostring Technologies) for 30 min at RT in a humidity chamber. Next, 500 nM Syto83 and antibodies targeting CD4 (clone EPR6855, Abcam), CD20 (clone IGEL/773, Novus) in Buffer W were applied to each section for 1 hour at RT. Slides were washed twice in fresh 2X SSC then loaded on the GeoMx Digital Spatial Profiler (DSP).

Entire slides were imaged at 20X magnification and morphologic markers were used to select 12 Region Of Interest (ROI) either using circle or organic shapes. ROIs were exposed to 385 nm light (UV), releasing the indexing oligonucleotides which were collected with a microcapillary and deposited in a 96-well plate for subsequent processing. The indexing oligonucleotides were dried down overnight and resuspended in 10 μl of DEPC-treated water.

Sequencing libraries were generated by PCR from the photo-released indexing oligos and AOI-specific Illumina adapter sequences, and unique i5 and i7 sample indices were added. Each PCR reaction used 4 μl of indexing oligonucleotides, 4 μl of indexing PCR primers, 2 μl of Nanostring 5X PCR Master Mix.

Thermocycling conditions were 37°C for 30 min, 50°C for 10 min, 95°C for 3 min; 18 cycles of 95°C for 15 s, 65°C for 1 min, 68°C for 30 s; and 68°C for 5 min. PCR reactions were pooled and purified twice using AMPure XP beads (Beckman Coulter), according to the manufacturer’s protocol. Pooled libraries were single-sequenced at 27 base pairs and with the single-index workflow on an Illumina NovaSeq S4 instrument. Novaseq-derived FASTQ files for each sample were compiled for each compartment using the bcl2fastq program of Illumina, and then demultiplexed and converted to digital count conversion (DCC) files using the GeoMx DnD pipeline version 2.0.0.16 of Nanostring according to manufacturer’s pipeline. DCC files were imported back into the GeoMx DSP instrument for QC and data analyses using GeoMx DSP analysis suite version 2.4.2.2 (Nanostring). QC was performed using the R package, GeoMxTools v3.4.0. Probes were checked for outlier status. A probe is removed if the geometric mean of that probe’s counts from all segments divided by the geometric mean of all probe counts representing the target from all segments is less than 0.1 or if the probe is an outlier according to the Grubb’s test in at least 20% of segments. The counts for all remaining probes for a given target were then collapsed into a single metric by taking the geometric mean of probe counts. Segments and genes were filtered out with abnormal low signal. Segments and genes with less than 5% of gene or segments detected and below the limit of quantification (LOQ) set at 2 geometric standard deviation above the geometric mean were removed from the study. Gene counts were normalized to the geometric mean of the 75^th^ percentile across all AOIs to give the upper quartile (Q3) normalization factors for each AOI.

### Cytokine quantification

Supernatants from melanoma tumor cell lines were analysed using Biolegend’s LEGENDplex^TM^ bead- based immunoassays to quantify IL-6, FLT3L, GM-CSF, IL-3, IL-34, IL-11, SCF, LIF, CXCL12, IL-15, M-CSF, IL-7 (Human Hematopoietic Stem Cell Panel) and IL-17A, CCL4, TNF-a, IL-4, IL-10, CCL5, sFas, sFasL, IFN-g, GZMA, GZMB, PRF1, GNLY (Custom Panel). Briefly, antibodies specific for the analytes were conjugated to different fluorescence-encoded beads. The beads were mixed with the supernatants, incubated for 2 hours at RT, washed, and incubated for 1 hour with detection antibodies. Finally, streptavidin-PE was added and incubated for 30 min, and the beads were washed and acquired using Attune NxT (Thermo Fisher Scientific). The results were analysed using LEGENDplex^TM^ Software Suite.

### Tumor conditioned medium production and T cell exposure

Melanoma cell lines (T672E, Me215, T333A, Me252, T311B, T618A) were seeded in 1mL of RPMI 10% FCS in 12-well plate, the day after medium was discarded, cells were washed with DPBS 1X and 2mL of fresh medium was added. On the third day the supernatant was collected, centrifuged to remove residual cells and frozen for further use. The day of the experiment KLRG1^+^ CD4 T cells were cultured in 100 ml of tumor conditioned medium and 100 ml of RPMI 8% HS with 100 IU/ml of IL-2 for 48 hours, then RT-qPCR was performed as described.

### Statistical analysis and illustrations

Statistical analyses were performed as indicated in the legend for each figure using GraphPad Prism (version 10). For all analyses, a p < 0.05 was statistically significant. Not statistically significant differences were left unlabelled. Illustrations were created with BioRender.com.

### Bioinformatic analyses

#### Single-cell RNA-seq data processing

For scRNA-seq data, Cell Ranger (version 6.1.1; 10x Genomics) was used to process and demultiplex raw sequencing data. Raw basecall files were converted to FASTQ format and aligned to the human reference genome (GRCh38) to generate single-cell feature counts. Raw gene-expression matrices were imported into R (v4.2.0), and Seurat was used for processing the datasets [50]. Feature-barcode hashing counts were stored as an additional assay and hashtag feature names were standardized per capture. Quality control was performed on each sample by computing mitochondrial content and excluding low-quality cells with fewer than 300 detected genes or mitochondrial content above 8%. Hashtag data were normalized with centred log-ratio transformation and demultiplexed with a MULTIseq approach [51] using automatic thresholds finding to define the best quantile; cells classified as doublets or negatives were removed, retaining only singlets. Sample metadata (patient, HLA, specificity, and tissue of origin) were assigned based on hashtag identity.

#### Cell type annotation

Datasets from multiple captures were integrated using STACAS [52] to mitigate technical effects while preserving biological variation. Unsupervised graph-based clustering was performed on the integrated total CD4^+^ T-cell dataset (PCA-based neighborhood graph followed by community detection) and visualized with UMAP. Cluster identities were assigned using canonical marker genes. To validate and harmonize these data-driven states, total CD4^+^ cells were mapped onto a human CD4 tumor-infiltrating T-cell reference atlas using ProjecTILs [51] (CD4T_human_ref_v2) [Garnica, Josep (2023). ProjecTILs human reference atlas of CD4^+^ tumor-infiltrating T cells, version 2. figshare. Dataset. https://doi.org/10.6084/m9.figshare.24886611.v1]. The resulting annotated total CD4^+^ embedding was then used as the reference framework to project tumor-reactive CD4^+^ T cells with ProjecTILs [51], enabling assignment of aligned functional clusters in the context of the total CD4 landscape.

#### Public dataset analysis

A pan-cancer atlas of T cells dataset [53] was retrieved as a pre-processed Seurat object. T cells were classified into CD4^+^ and CD8^+^ subsets based on marker gene expression (*CD4* and *CD8A/B*). For validation and cross-platform annotation consistency, CD4^+^ cells were projected onto the ProjecTILs CD4 reference map (CD4T_human_ref_v2 [Garnica, Josep (2023). ProjecTILs human reference atlas of CD4^+^ tumor-infiltrating T cells, version 2. figshare. Dataset. https://doi.org/10.6084/m9.figshare.24886611.v1]) using sample-level stratification.

#### Cell type gene markers

Marker genes for the cytotoxic CD4^+^ T-cell state were identified in the total CD4^+^ T-cell dataset by differential expression analysis comparing the Cytotoxic cluster against all other total CD4^+^ cells using Seurat’s FindMarkers function [47] on the RNA assay (test = “bimod”). Genes were required to be detected in at least 40% of cells in the Cytotoxic cluster, show a minimum difference in detection rate of 20% relative to other cells, and be expressed in at least 10 cells.

#### Pseudobulk differential expression analysis

To compare transcriptional profiles between KLRG1^+^ and KLRG1^−^ CD4^+^ T cells in peripheral blood (PBMC), pseudobulk differential expression analysis was performed. Cells from PBMC samples were stratified by KLRG1 expression status (defined by scGate purity classification [50]) and patient identity. Only patients with more than 10 cells per condition were retained for analysis. Gene expression raw counts were aggregated at the patient and KLRG1 expression level. T-cell receptor (TCR) genes were excluded from the analysis to avoid patient-specific confounding effects. A DESeq2 linear model [54] was fitted with patient as a covariate to account for inter-patient variability and KLRG1 expression as the variable of interest (design: ∼ Patient + KLRG1_expression). Genes with total counts below 20 across all samples were filtered out. Differential expression was assessed using the Wald test, followed by log2 fold-change shrinkage using the apeglm method [55] to reduce high-variance estimates.

#### Gene Ontology enrichment analysis

Gene Ontology (GO) enrichment analysis was performed using the clusterProfiler package [56]. Gene sets were obtained from the Molecular Signatures Database (MSigDB) via the msigdbr R package [Dolgalev; https://igordot.github.io/msigdbr/], restricting to the C5 category (Gene Ontology gene sets) and specifically to biological processes (GOBP). Differentially expressed genes from the pseudobulk analysis (upregulated: log2FC > 0, adjusted p-value < 0.05) were tested for over-representation against a background of all expressed genes using Fisher’s exact test, with false discovery rate (FDR) correction via the Benjamini-Hochberg method. Significantly enriched GO terms were defined as having adjusted p-value < 0.05.

#### Protein–protein interaction network analysis

For protein–protein interaction network analysis centred on KLRG1, adhesion-related GO terms were identified from the enrichment results by filtering for terms containing “adhesion,” “cell adhesion,” or “cell-cell” in their descriptions. Genes contributing to these adhesion-related GO terms and upregulated in KLRG1^+^ CD4^+^ T cells were extracted and combined with KLRG1 to define the gene set for network analysis. Protein–protein interactions were queried from the STRING database (version 11.5, Homo sapiens, confidence score threshold = 400) using the STRINGdb R package [57]. Gene symbols were mapped to STRING identifiers, and only interactions directly involving KLRG1 (first-degree interactions) were retained to generate a KLRG1-centered network.

#### Spatial transcriptomics processing

GeoMx Digital Spatial Profiler (DSP) RNA whole transcriptome atlas (WTA) data were analysed using the NanoString GeoMx NGS Pipeline and GeoMxTools R package (doi:10.18129/B9.bioc.GeomxTools). Raw DCC files containing sequencing counts were imported along with probe annotation files to map probe identifiers to gene symbols. Regions of interest (ROIs) were annotated based on tissue morphology and marker expression: CD4^+^ T cell regions (CD4^+^ enriched), CD20^+^ B cell regions (CD20^+^ enriched). Additional spatial metadata included tumor localization (inside vs outside tumor), B cell distance categories (close vs far) for CD4^+^ regions, and patient identity. Raw count data from different ROI types were merged, with genes absent in specific assays assigned zero counts. Quality control included assessment of total read counts and size factor distribution per ROI. Data normalization was performed using DESeq2 [54] size factor estimation followed by variance-stabilizing transformation (VST) to account for sequencing depth differences and stabilize variance across the dynamic range of expression.

#### Spatial transcriptomics deconvolution

Cell-type deconvolution was performed using UCell [56], a rank-based signature scoring method that quantifies gene-set enrichment from expression profiles. Cell-type-specific gene signatures were derived from the manually annotated CD4^+^ T-cell functional clusters using only TILN cells (to match the tissue context of the spatial dataset). For each cluster, marker genes were identified by differential expression analysis (bimod test), retaining genes with adjusted p-value < 0.05, log2 fold change > 0.4, and sufficient prevalence (detected in at least 40% of cells), while enforcing a minimum difference in detection rate of 10% versus other clusters and excluding non-informative genes (HLA and ribosomal genes). The top 10 ranked marker genes per cluster were used to define the final signatures. UCell enrichment scores were then computed for each signature across all GeoMx ROIs using the raw count matrices, yielding quantitative scores that reflect the relative enrichment of each CD4^+^ T-cell functional state within each spatial region.

#### Spatial transcriptomics differential expression analysis

Differential expression analyses across ROI groups were performed using DESeq2 [52] by fitting negative binomial generalized linear models with a design formula including the factor of interest. Log2 fold-change shrinkage was applied using the apeglm method [53] to stabilize high-variance estimates.

#### Data and code availability

The single-cell RNA sequencing (scRNA-seq) data generated in this study have been deposited in Zenodo and are publicly available under the DOI: https://doi.org/10.5281/zenodo.18683366. GeoMX data is publicly available under the DOI: https://doi.org/10.5281/zenodo.18761515.

All bioinformatics analyses used to generate the results and figures in this study are available in a public GitHub repository: https://github.com/carmonalab/KLRG1_CD4-CTL_reproducibility.

## Acknowledgments

We are grateful to the patients for their dedicated collaboration, and to the healthy donors for their blood donations. We thank the FACS Facility, Genomics Platform, Protein and Peptide Facility at the University of Geneva and the Immune Landscape Laboratory Platform at University of Lausanne for their collaboration in this work and the Ludwig Institute for Cancer Research. We acknowledge Oscar Vadas and Rémy Visentin at the Protein Biochemistry Platform of the University of Geneva for assistance with SpCas9 production. We also thank the funding sources that supported this work: M.C. is the recipient of an iGE3 PhD Salary Award (University of Geneva), Gabbiani Fund Award (University of Geneva) and EFIS-IL Short-term Fellowship. M.C. and R.C. are supported by NIHR Oxford Biomedical Research Centre (NIHR203311) and Oxford NIHR Blood and Transplant Research Unit (BTRU) in Precision Cellular Therapeutics (NIHR203339). V.H. and B.W. were attributed the ISREC TANDEM grant and B.W. a Gilead grant (25481). C.J. is the recipient of a SNSF PRIMA fellowship PR00P3-179727, Fondazione San Salvatore fellowship and an ISREC TANDEM grant.

## RESULTS

### Single-cell profiling of melanoma patients identifies a recurrent cytotoxic CD4 T cell state enriched in peripheral blood and present among tumor-specific T cells

Using a high-throughput single-cell cytotoxicity assay in picowells, we previously demonstrated delayed tumor cell killing by human tumor-specific CD4 compared with CD8 T cells [14]. However, the molecular mechanisms underlying this phenomenon were not explored. Analysis of the picowell array dataset (Fig. 1A, left) revealed that cytotoxic CD4 T cells, in picowells where tumor killing occurred, engaged in more frequent but shorter-duration contacts with their targets compared with CD8 T cells (Fig. 1A, right). This behaviour was not due to defective immune synapsis (IS) formation, as CD4 T cells efficiently polarized both centrosomes (Fig. 1B, top) and lytic granules (Fig. 1B, bottom) towards the IS upon contact with their targets, as confirmed by high-resolution cryo-expansion microscopy [41, 42].

**FIGURE 1.**
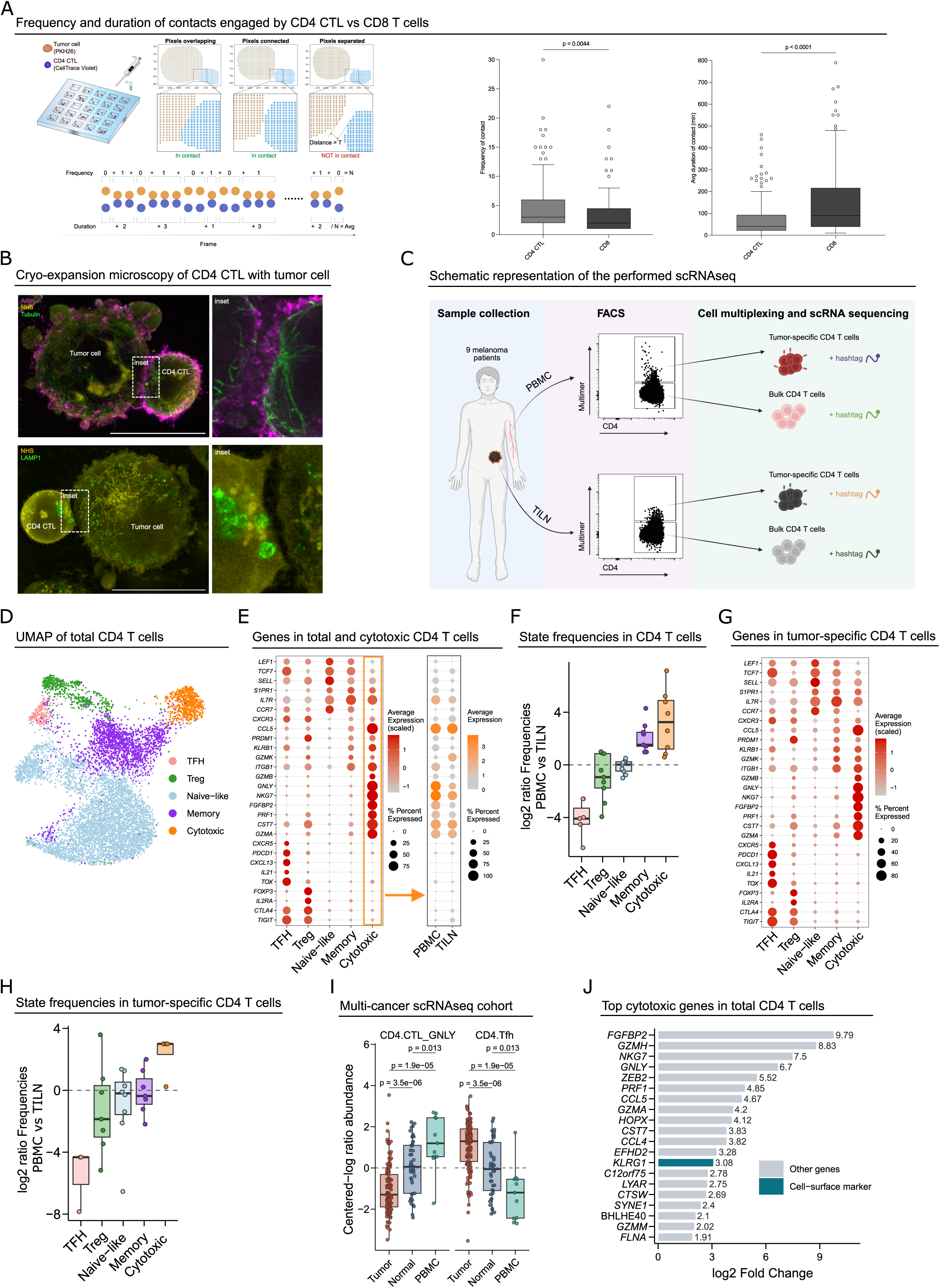
Killing dynamics and transcriptional states of tumor-specific cytotoxic CD4 T cells. (A) Schematic experimental representation (left). Frequency and duration of contacts engaged by CD4 CTL compared to CD8 T cells (right). Data are represented using Tukey box and whiskers plot with median (in-box line) and outliers (dots), Mann-Whitney unpaired test. (B) Cryo-expansion microscopy of CD4 CTL-tumor cell pairs labelled for, top panel, Actin (magenta), NHS (yellow), Tubulin (green) or, bottom panel, for Lamp 1 (green). Insets show magnifications of the immune synapse, scale: 10 µm. (C) Schematic representation of the experimental overview of the performed scRNA-seq. (D) UMAP of the integrated total CD4^+^ T-cell scRNA-seq dataset, colored by manually curated cell-state annotation (E) Dot plots of canonical marker genes across total CD4^+^ states (left) and within the Cytotoxic state stratified by tissue (PBMC vs TILN; right). Dot size indicates the fraction of expressing cells and color indicates average expression (scaled per gene in the left panel; average normalized expression in the Cytotoxic PBMC vs TILN panel). (F) Per-patient log2 ratio of state frequencies between PBMC and TILN for total CD4+ cells. Boxes summarize patients; dashed line indicates equal abundance. (G) Tumor antigen–specific CD4^+^ T cells annotated based on the total CD4 dataset shown in (D); dot plot shows marker expression across projected CD4^+^ states. (H) Per-patient log2 ratio of projected functional-cluster frequencies between PBMC and TILN within the tumor antigen–specific CD4^+^ compartment. (I) Multi-cancer scRNA-seq cohort: centered log-ratio abundance of CD4.CTL_GNLY and CD4.Tfh across tissues (Tumor, Normal, PBMC). Points represent samples; adjusted P values are from BH-corrected Wilcoxon tests. (J) Top 20 differentially expressed marker genes of the Cytotoxic CD4^+^ state versus all other total CD4^+^ cells within total CD4^+^ T cell dataset. Bars show log2 fold change. Genes are grouped as predicted cell-surface receptors versus other genes based on Gene Ontology annotations.

To define cytotoxic gene programs in human tumor-specific CD4 T cells, we performed single-cell RNA sequencing on samples obtained from nine melanoma patients (Fig. 1C). Total CD4 T cells, together with CD4 T cells specific for the tumor antigens NY-ESO-1_87-99_, NY-ESO-1_123-137_ and MAGE-A3_243-258_, were sorted using fluorescent peptide-MHC class II multimers from peripheral blood mononuclear cells (PBMC) and paired tumor-infiltrating lymph nodes (TILN) (Supp. Fig.1A).

Unsupervised clustering of single-cell RNA-seq profiles from total CD4 T cells identified seven transcriptionally distinct clusters (Supp. Fig.1B). Based on canonical marker expression, these clusters were annotated into five major CD4 T cell states: Naïve-like, Memory, T follicular helper (Tfh), T regulatory (Treg), and cytotoxic CD4 T cells (CD4.CTL) (Fig. 1D). Naïve-like cells expressed classical markers including *LEF1*, *TCF7*, *SELL*, *CCR7*, and *IL7R*, whereas Memory cells were characterized by *CXCR3*, *CCL5*, *PRDM1*, and *ITGB1*. The Tfh cluster expressed *CXCR5*, *PDCD1*, *TOX*, and *IL21*, while Treg cells were defined by *FOXP3*, *IL2RA*, and *CTLA4*. A distinct cytotoxic CD4 T cell cluster was defined by high expression of cytotoxic effector genes, including *GZMB*, *GNLY*, *PRF1*, *NKG7*, and *FGFBP2* (Fig. 1E; Supp. Fig.1C). Cell type annotations were independently validated by mapping cells to a curated human CD4 T cell reference atlas (Supp. Fig.1D).

We next examined the distribution of these CD4 T cell states across PBMC and TILN. Cytotoxic CD4 T cells were consistently enriched in PBMC compared with TILN across patients, with a median PBMC/TILN fold change of 10.5. In contrast, Tfh cells showed the opposite pattern and were strongly enriched in TILN (median fold change 0.05) (Fig. 1F). Moreover, the relatively small number of cytotoxic CD4 T cells detected in TILN exhibited a reduced cytotoxic transcriptional signature compared with their PBMC counterparts (Fig. 1E; Supp Fig.1E).

CD4.CTLs expressing the cytotoxic gene module were also detected among tumor antigen–specific CD4 T cells, indicating that cytotoxic CD4 T cells can arise within tumor-reactive responses (Fig. 1G; Supp Fig.1F). Notably, the same tissue distribution pattern was observed within the antigen-specific compartment, with cytotoxic CD4 T cells preferentially detected in PBMC, whereas Tfh cells were enriched in TILN (Fig. 1H).

To assess generalisability, we analysed publicly available single-cell RNA-seq datasets of T cells from peripheral blood, tumors, and normal adjacent tissues across multiple cancer types [49]. An effector CD4.CTL population, defined by expression of the cytotoxic program (*GZMB*, *GNLY*, *PRF1*, *NKG7*, and *FGFBP2*), was recurrent across diverse malignancies, alongside Tfh cells (Supp Fig.1G). Consistent with our cohort, CD4.CTLs were enriched in peripheral blood relative to tumors, whereas Tfh cells were preferentially enriched within tumors, with intermediate frequencies observed in non-malignant solid tissues (Fig. 1I).

To identify candidate surface markers of cytotoxic CD4 T cells, we performed differential expression analysis between cytotoxic CD4 T cells and other CD4 T cell populations. Both in our dataset (Fig. 1J) and across multiple cancer cohorts (Supp. Fig.1H), the Killer Cell Lectin-like Receptor G1 (*KLRG1*) was the most differentially expressed gene encoding a cell surface marker in the CD4.CTL population.

Together, these data identify a cytotoxic CD4 T cell program enriched in peripheral blood that is present among tumor antigen-reactive cells and includes KLRG1 as a potential surface marker.

### KLRG1 marks functionally active circulating human cytotoxic CD4 T cells, with *GNLY* and *GZMB* as key cytotoxic molecules

To determine whether KLRG1 marks cytotoxic CD4 T cells, we stratified total CD4 T cells by *KLRG1* expression and quantified the relative abundance of cytotoxic versus non-cytotoxic states within each compartment in the scRNA-seq data. KLRG1^+^ cells were enriched for the CD4.CTL state, whereas KLRG1^−^ cells were predominantly composed of non-cytotoxic CD4 populations (Fig. 2A, left). This shift in composition was recapitulated in the independent multi-cancer single-cell cohort (Fig. 2A, right). Differential expression analysis comparing KLRG1^+^ versus KLRG1^−^ CD4 T cells revealed upregulation of a canonical cytotoxic program in KLRG1^+^ cells, including *GNLY*, *PRF1*, *NKG7*, *CST7,* and multiple granzymes (*GZMA, GZMB*, *GZMK*, and *GZMH*) (Fig. 2B).

**FIGURE 2.**
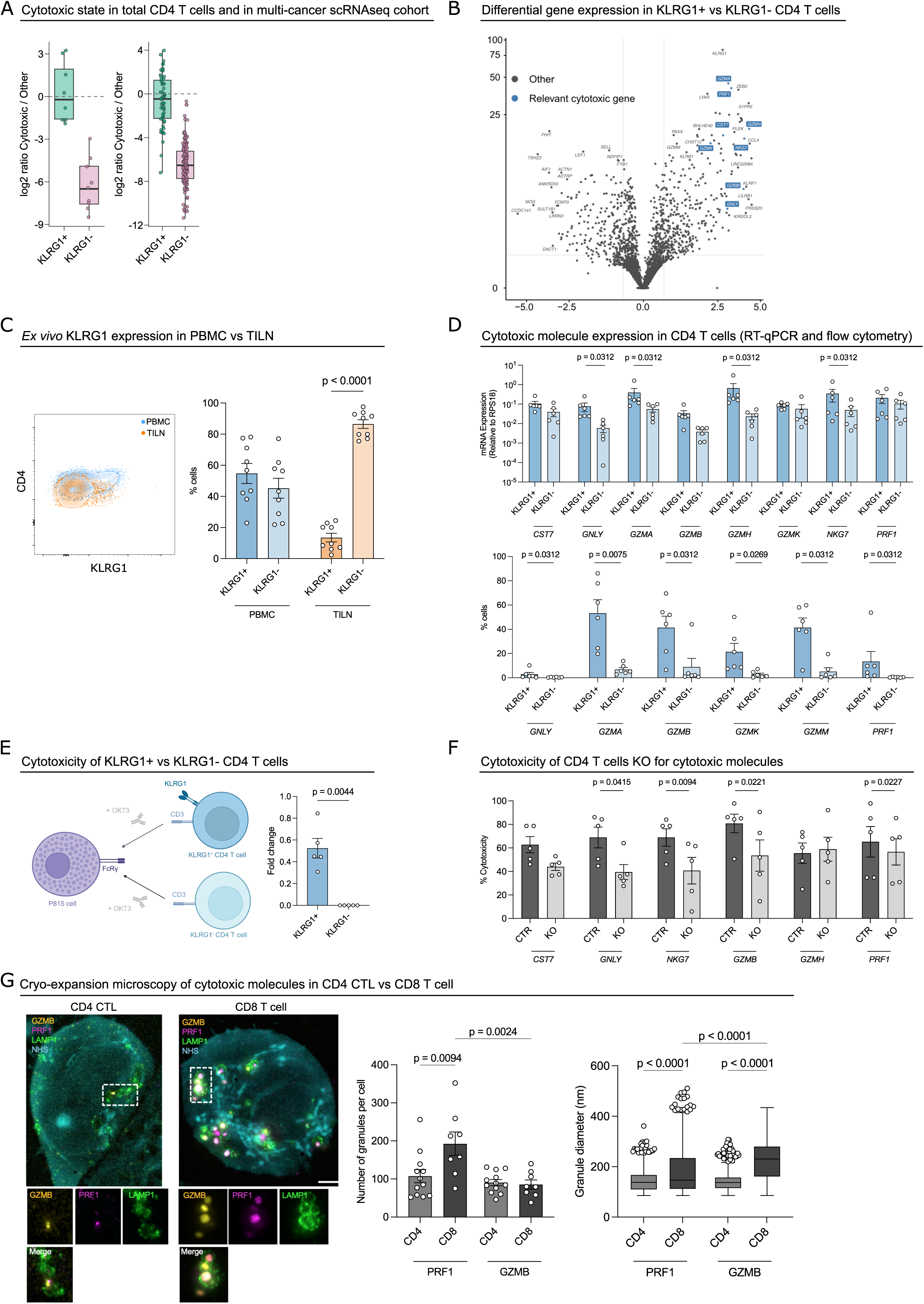
KLRG1 marks cytotoxic CD4 T cells and modulates effector function. (A) Log2 ratio of Cytotoxic versus non-cytotoxic CD4^+^ T cells within KLRG1^−^ and KLRG1^+^ compartments in the total CD4^+^ dataset (left) and an independent multi-cancer single-cell cohort (right). Boxes indicate median and interquartile range; points represent individual samples. (B) Volcano plot of differential expression between KLRG1^+^ and KLRG1− PBMC-derived CD4^+^ T cells in the total CD4^+^ dataset, performed by pseudobulk to account for inter-sample variation. Cytotoxic-associated genes are highlighted in blue; other genes are shown in grey. Dashed lines indicate fold-change and adjusted p-value thresholds. (C) Flow cytometry dot plot of a representative example of KLRG1^+^ *ex vivo* expression in PBMC vs TILN and cumulative results represented in the graph below. The data are shown as the means ± standard errors of the mean (SEMs), 2way ANOVA. (D) Cytotoxic molecule expression in CD4 T cells in PBMC measured by RT-qPCR (top) (n=6) and flow cytometry (bottom) (n=6). Data are shown as the means ± SEMs, Wilcoxon paired t-test. (E) LDH-based cytotoxicity assay of *ex vivo* sorted KLRG1^+^ CD4 T cells, with scheme (left) and data (right). The data are shown as the means ± SEMs, Wilcoxon paired t-test (n=5). (F) LDH-based cytotoxicity assay of the knock-out (KO) vs wild-type (WT) *CST7*, *GNLY*, *NKG7*, *GZMB*, *GZMH, PRF1* by CRISPR/Cas9, paired t-test (n=5). (G) Cryo-expansion microscopy of CD4 and CD8 T cells labelled for GZMB (yellow), PRF1 (magenta), LAMP1 (green) and NHS (blue), scale: 1 µm. On the right, quantification of granules per cell, one-way ANOVA, and granule diameter represented using Tukey box and whiskers plot with median (in-box line) and outliers (dots), Kruskal-Wallis unpaired test.

At the protein level, CD4 T cells were stained with an anti-KLRG1 antibody, revealing that 54% of circulating CD4 T cells in patient blood expressed KLRG1. In contrast, tumor-infiltrating CD4 T cells were predominantly KLRG1^−^ (Fig. 2C).

We next sorted KLRG1^+^ and KLRG1^-^ CD4 T cell populations from the peripheral blood of healthy donors and performed mRNA expression analyses of the cytotoxic genes identified in the scRNA-seq data (Fig. 2B). These analyses confirmed that KLRG1^+^ CD4 T cells expressed significantly higher levels of cytotoxic genes, particularly *GNLY, GZMA, GZMB,* and *GZMH*, compared with their KLRG1^−^ counterparts (Fig. 2D, top).

To further assess protein expression, CD4 T cells were co-stained for these markers together with KLRG1. In PBMC-derived CD4 T cells, high KLRG1 expression was associated with elevated levels of GZMK, GZMA, GZMM and GZMB (Fig. 2D, bottom). The enhanced cytotoxic potential of KLRG1^+^ cells was functionally confirmed using a redirected cytotoxicity assay (Fig. 2E, left), which demonstrated their superior ability to kill target cells compared with KLRG1^-^ counterparts (Fig. 2E, right). This was accompanied by a trend toward increased secretion of pro-inflammatory and cytotoxic mediators, including TNF-α, CCL4, CCL5, IFN-γ and GNLY (Supp Fig.2A).

To identify key mediators of CD4 T cell cytotoxicity, we focused on six effector genes differentially expressed in CD4.CTLs: *CST7*, *GNLY*, *NKG7*, *GZMB*, *GZMH,* and *PRF1*. Using CRISPR/Cas9- mediated gene editing, we individually knocked out these genes in cytotoxic tumor-antigen specific CD4 T cell clones derived from primary human CD4 T cells isolated from patients’ samples. Deletion of *GNLY*,

*NKG7*, *GZMB* or *PRF1* resulted in a significant reduction in cytotoxic activity, demonstrating their critical role in mediating CD4 CTL function (Fig. 2F). in contrast, deletion of CST7 or *GZMH* had no measurable effect, suggesting that these genes are dispensable for cytotoxic function.

To further characterize the effector machinery at the protein and subcellular levels, we performed cryo– expansion microscopy. This revealed quantitative differences in lytic granule features between CD4 CTLs and CD8 T cells: CD4 CTLs displayed reduced PRF1-associated granule content and smaller GZMB-positive lytic structures compared with CD8 T cells (Fig. 2G). Consistently, imaging of GNLY together with the lysosomal marker LAMP1 confirmed GNLY localization to cytotoxic granules and highlighted cell-type-specific differences in GNLY granule abundance and morphology (Supp Fig.2B). Together, these transcriptional, genetic, functional, and imaging data demonstrate that circulating KLRG1^+^ CD4 CTLs deploy a bona fide cytotoxic program.

### CD325 expression is essential for KLRG1^+^ CD4 T cell cytotoxic activity

To define mechanisms underlying tumor cell elimination by cytotoxic CD4 T cells, we performed pathway enrichment analysis of genes upregulated in KLRG1⁺ CD4 T cells compared with the KLRG1⁻ fraction in PBMC samples. Among the most significantly enriched pathways, “cell–cell adhesion” emerged as a top hit (Fig. 3A), suggesting that adhesive interactions contribute to cytotoxic function of KLRG1⁺ CD4 T cells (Supp Fig.3A).

**FIGURE 3.**
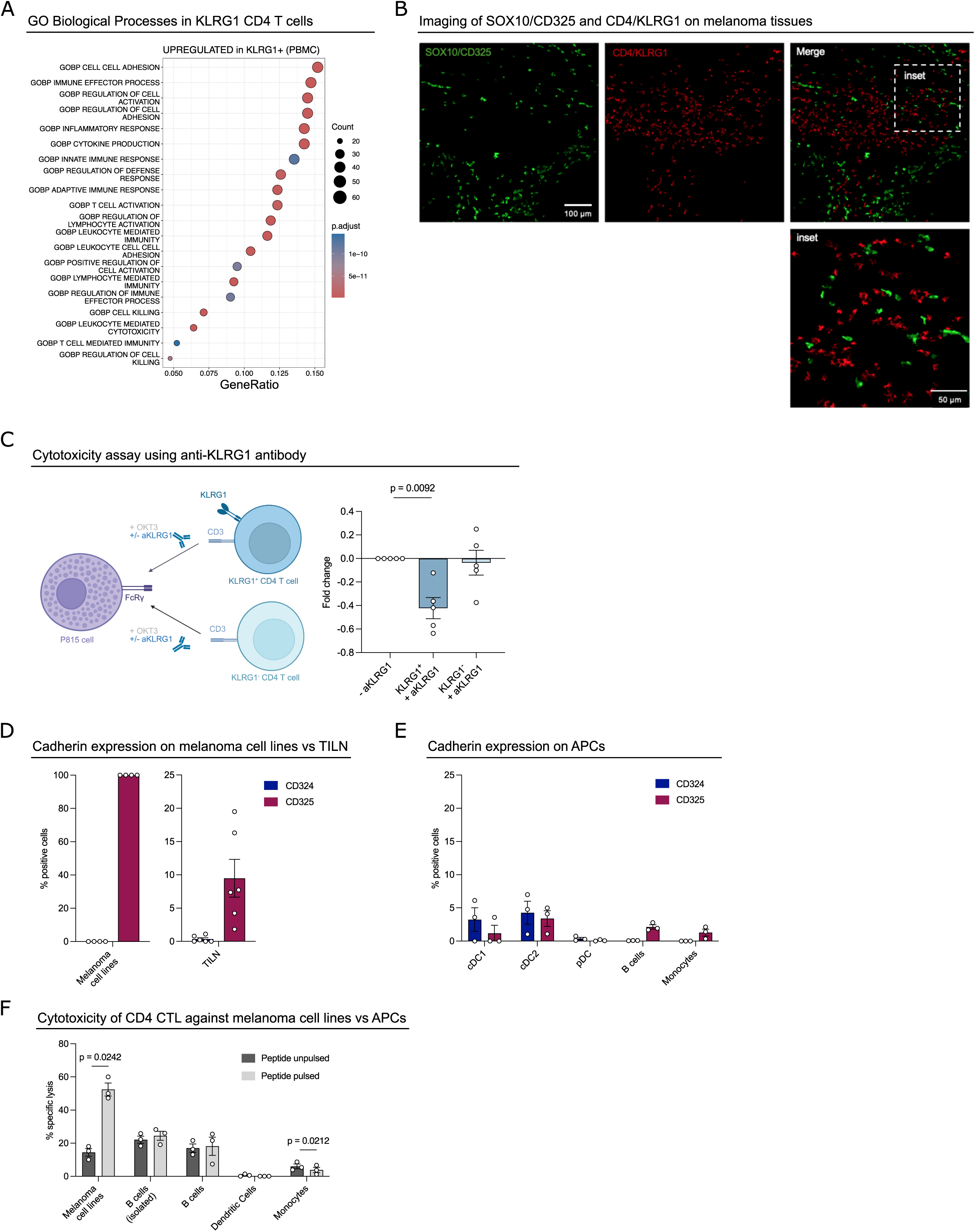
Tumor-specific KLRG1 ligands. (A) Gene Ontology (GO) Biological Process enrichment analysis of genes upregulated in KLRG1^+^ versus KLRG1− CD4^+^ T cells in PBMC (derived from the differential expression in Fig. 2B). The top 20 enriched GO terms are shown, ranked by GeneRatio; dot size indicates the number of genes contributing to each term and dot color indicates the adjusted p-value. (B) Multispectral imaging assessment of SOX10/CD325 and CD4/KLRG1 expression directly in situ on melanoma tissues. The insets illustrate high-power images. (C) LDH-based cytotoxicity assay using anti-KLRG1 antibody scheme (left) and data (right), paired t-test. (D) Expression of CD324 and CD325 on melanoma cell lines (n=4) and on tumor cells from TILN (n=6). (E) Expression of CD324 and CD325 on dendritic cells, B cells and monocytes (n=3). (F) Percentage of 7-AAD positive cells (peptide unpulsed vs pulsed) following co-culture with HLA-matched antigen-specific cytotoxic CD4 T cells, paired t-test (n=3).

KLRG1 ligands include E-cadherin (CD324) and N-cadherin (CD325) [58]. Immunohistochemical staining of melanoma tissue sections revealed that KLRG1⁺ CD4 T cells localise in proximity to tumor cells expressing CD325 (Fig. 3B), consistent with a functional interaction between KLRG1 and its ligand within the tumor microenvironment.

To determine whether the cytotoxic function of CD4 T cells depends on the expression of KLRG1 ligands on target cells, we performed killing assays in the presence of an anti-KLRG1 blocking antibody (Fig. 3C). A significant reduction in cytotoxicity was observed when KLRG1^+^ CD4 T cells were treated with the blocking antibody, whereas no effect was observed in KLRG1^-^ CD4 T cells.

MHC class II is typically expressed on professional APCs, raising the possibility that these cells could be targeted by cytotoxic CD4 T cells, as recently observed in mouse models [59]. CD324 expression was minimal in melanoma cell lines and TILN, whereas melanoma cell lines were uniformly CD325^+^, and approximately 10% of tumor cells infiltrating lymph nodes expressed CD325 (Fig. 3D and Supp Fig.3B), in line with the spatial analyses (Fig. 3B). Both KLRG1 ligands, CD324 and CD325, were absent from human APCs, including B cells, monocytes, cDC1, cDC2 and pDC subsets (Fig. 3E and Supp Fig.3C).

Consistent with this, when co-cultured with cytotoxic CD4 T cells, HLA-matched, peptide-pulsed APCs were resistant to killing, in contrast to melanoma cells, which were efficiently lysed (Fig. 3F). These data demonstrate that the absence of KLRG1-cadherin interactions protects APCs from CD4 T cell-mediated lysis, while rendering tumor cells selectively susceptible to killing.

### Cytotoxic CD4 T cells undergo functional reprogramming within the tumor microenvironment

scRNA-seq analysis revealed a compartmentalization of CD4 T cell states, with cytotoxic CD4 T cells enriched in peripheral blood and tumor samples dominated by Tfh cells (Fig. 1F, H, I). To validate this shift, we performed RT–qPCR on sorted CD4 T cells from blood and TILN. Consistent with the single-cell data, TILN-derived CD4 T cells showed reduced expression of cytotoxic effector genes (*GNLY*, *GZMA*, *GZMB*, and *NKG7*) (Fig. 4A) and higher expression of Tfh-associated genes (*BCL6*, *ICOS*, and *PDCD1*) compared with their blood-derived counterparts (Fig. 4B).

**FIGURE 4.**
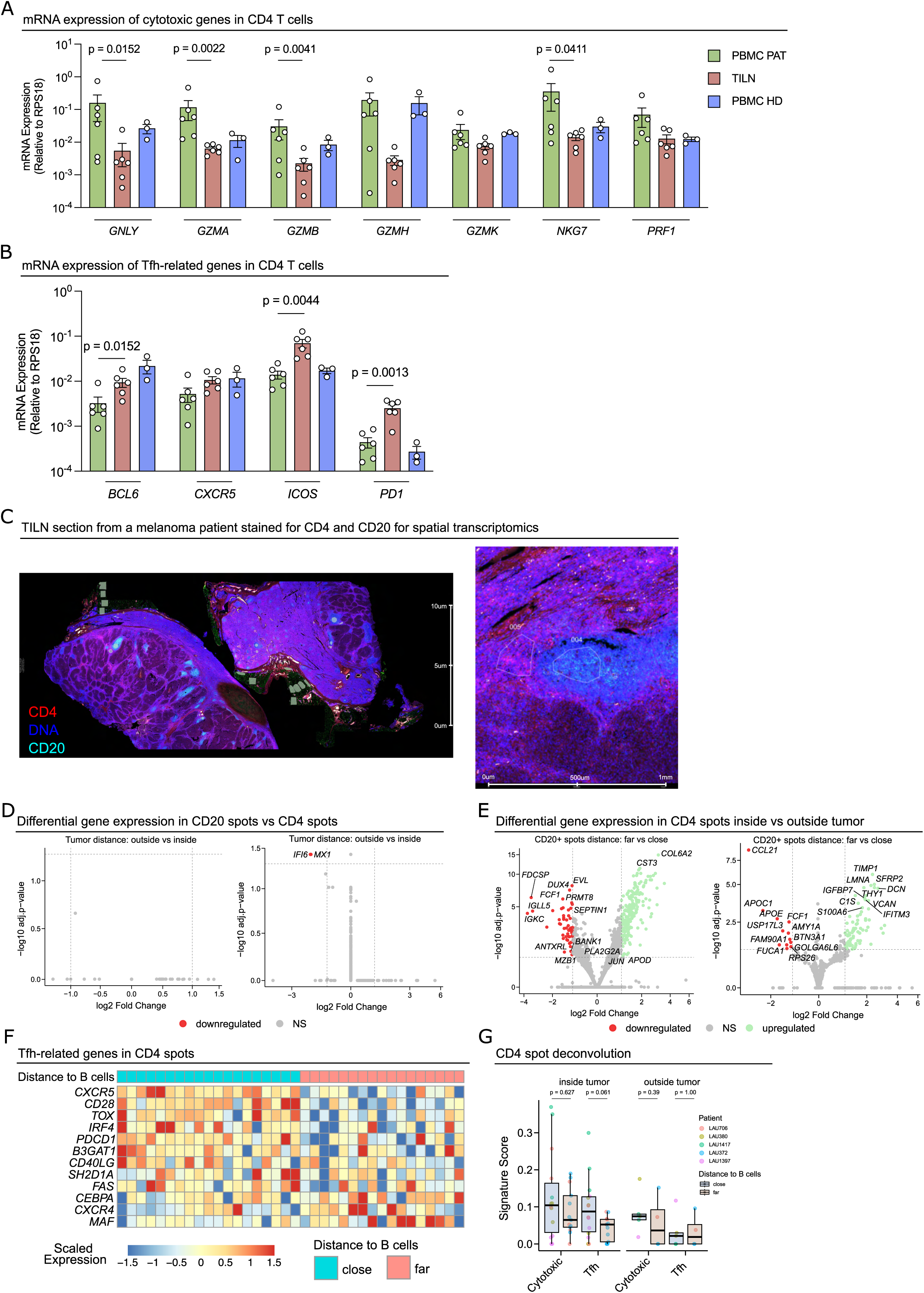
Spatial remodelling of cytotoxic CD4 T cells in the tumor microenvironment. (A) mRNA expression of cytotoxic genes in CD4 T cells from PBMC vs TILN vs HDs. The data are shown as the means ± SEMs, Mann-Whitney unpaired t-test (n=6). (B) mRNA expression of Tfh-related genes on CD4 T cells from PBMC vs TILN vs HDs. The data are shown as the means ± SEMs, Mann-Whitney unpaired t-test (n=6). (C) Representative example of a TILN section from a melanoma patient stained with CD4 in red and CD20 in blue (left), zoom of a selected region for CD4 T cells (005) and for B cells (004) (right). (D–F) Volcano plots showing differential gene expression in spatial ROIs. (D) Differential gene expression analysis of CD20⁺ (B cell–enriched, left) and CD4⁺ (right) ROIs comparing regions outside versus inside the tumor bed. (E) Differential gene expression in CD4⁺ ROIs according to proximity to B cells (far vs close), stratified by tumor localization: inside (left) and outside (right). (F) Heatmap of selected Tfh-associated genes across CD4^+^ spots. Expression values are displayed as row-wise scaled expression (z-score per gene). Columns represent individual CD4^+^ spots, annotated by proximity to B cells (close vs far). (G) Deconvolution-based signature scores (Cytotoxic and Tfh) for CD4^+^ spots, stratified by tumour localization (inside vs outside tumour) and distance to B cells (close vs far). Boxes show median and interquartile range; points represent individual spots colored by patient. P-values (Wilcoxon rank-sum test, BH-adjusted) compare close vs far within each localisation and signature.

To characterize the transition from CD4 CTLs to Tfh cells within the tumor microenvironment (TME), we performed spatial transcriptomics on tumor sections from five patients included in the scRNA-seq cohort (Fig. 4C). Given the role of Tfh cells in supporting B cell responses [60], we mapped CD4^+^- and CD20^+^- enriched regions of interest (ROIs), corresponding to CD4 T cells and B cells, respectively, to assess their spatial distribution both inside and outside the tumor bed. CD4 T cell-enriched regions were classified as proximal or distal to B cell ROIs (Supp Fig.4A).

CD20⁺ ROIs showed minimal transcriptional differences between regions inside and outside the tumor (Fig. 4D, left), whereas CD4⁺ ROIs within the tumor upregulated interferon-response genes, including *IFI6* and *MX1* (Fig. 4D, right). This expression pattern was consistent with our scRNA-seq analysis, which showed higher expression of these genes in TILN-derived CD4 T cells compared with PBMC (Supp Fig.4B). However, multiple genes were differentially expressed between CD4⁺ ROIs proximal to B cells and those distal from them. Immunoglobulin-related genes were enriched in CD4⁺ regions adjacent to B cells, consistent with the presence of B cells in these areas. These spatial transcriptional differences were most pronounced when CD4⁺ ROIs were located within the tumor (Fig. 4E).

CD4^+^ regions proximal to B cells were enriched for Tfh-associated markers, including *CXCR5*, *TOX*, and *PDCD1,* while showing reduced expression of *MAF, CXCR4, and CEBPA,* consistent with altered activation states (Fig. 4F).

To quantify CD4 T cell state across spatial ROIs, we computed rank-based signature scores for cytotoxic and Tfh programs in each CD4⁺ ROI using gene sets derived from our scRNA-seq–defined CD4 T cell states. When ROIs were stratified by proximity to B cells, cytotoxic scores did not differ between B-cell– proximal and distal regions, either inside or outside the tumor bed (Fig. 4G). In contrast, within the tumor, Tfh signature scores showed higher values in ROIs proximal to B cells (Fig. 4G), whereas outside the tumor bed, Tfh scores were comparable irrespective of B cell proximity.

Together, these data support a context-dependent association between proximity to B cells and the induction of Tfh-like programs, which is most evident within the tumor microenvironment.

### The tumor microenvironment modulates the cytotoxicity of CD4 T cells

To identify local factors within the TME that influence the cytotoxic phenotype of CD4 T cells, particularly in promoting a Tfh-like phenotype in proximity to B cells, we quantified soluble factors in the tumor-conditioned medium (CM) derived from six melanoma cell lines (Fig. 5A, left) and in supernatants from B cells isolated from patients’ TILN (Fig. 5A, right). IL-6, IL-11 and LIF, were enriched in tumor CM but not in B cell supernatants (Fig. 5A). To assess whether CD4 T cells respond to these factors, we examined expression of their receptors and the shared signalling subunit gp130.

**FIGURE 5.**
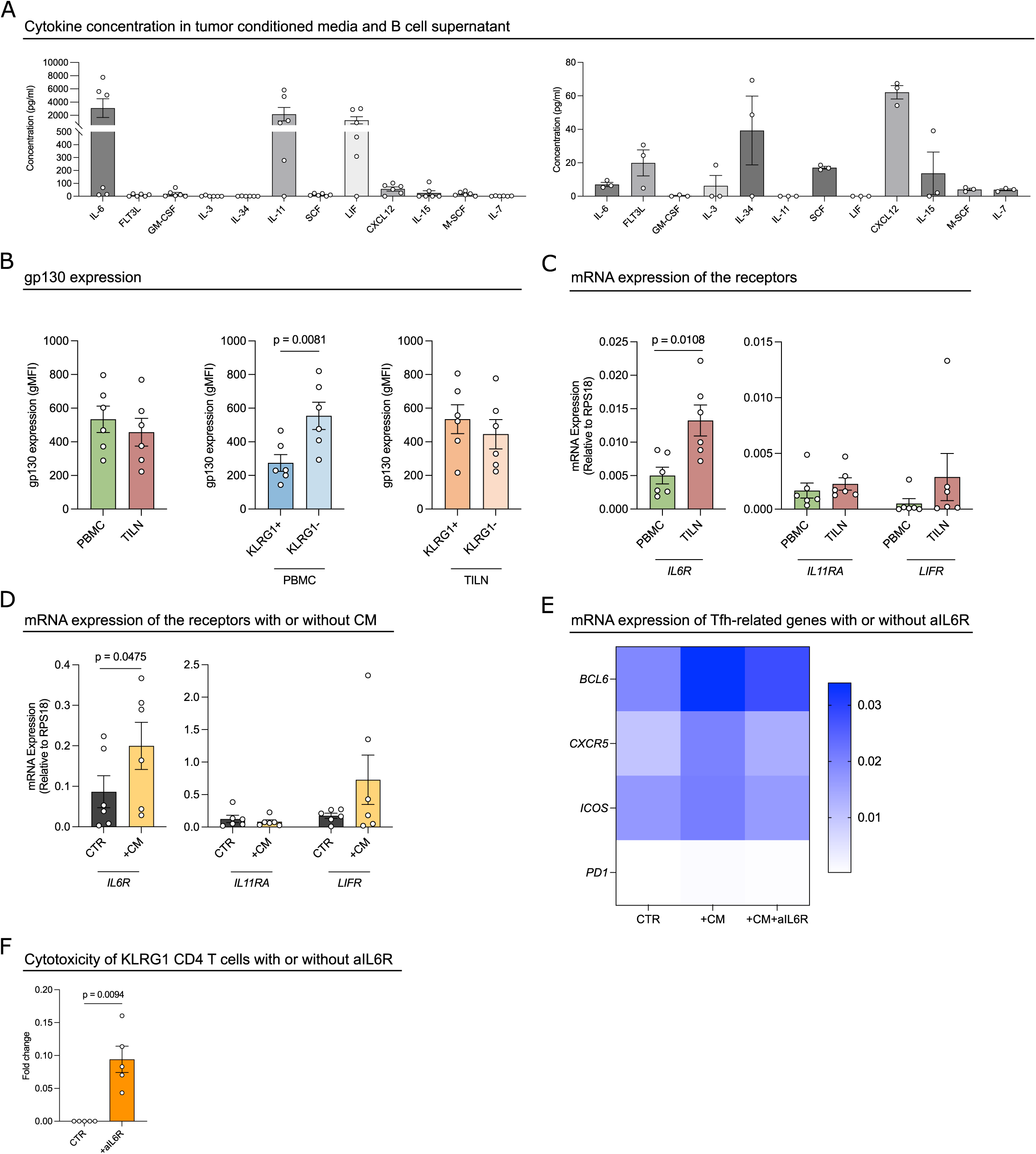
Tumor microenvironment cues drive phenotypic reprogramming of cytotoxic CD4 T cells. (A) Cytokine concentration (pg/ml) in tumor conditioned media (left) (n=6) and B cells supernatant (right) (n=3). (B) gp130 expression by flow cytometry in PBMC vs TILN, KLRG1^+^ vs KLRG1^-^ (n=6), Mann-Whitney unpaired t-test. (C) mRNA expression of the receptors *IL6R, IL11RA, LIFR* in PBMC vs TILN. The data are shown as the means ± SEMs, Mann-Whitney unpaired t-test (n=6). (D) mRNA expression of the receptors *IL6R, IL11RA, LIFR* in PBMC with or without addition of tumor conditioned medium (CM) on KLRG1^+^ CD4 T cells from PBMC, paired t-test (n=6). (E) mRNA expression of Tfh-related genes on KLRG1^+^ CD4 T cells with addition of tumor conditioned medium (CM), or with aIL-6R antibody compared to control (CTR). (F) Cytotoxicity assay of KLRG1^+^ CD4 T cells with or without addition of aIL6R antibody, paired t-test (n=5).

gp130 was expressed at similar protein levels in CD4 T cells from blood and tumor samples (Fig. 5B, left). However, in peripheral blood, gp130 expression was higher in KLRG1^-^ CD4 T cells compared with KLRG1^+^ CD4 T cells (Fig. 5B, middle), whereas this difference was lost within the tumor, where both populations exhibit reduced cytotoxicity (Fig. 5B, right).

IL-6 receptor (IL-6R) expression was significantly increased in TILN-derived CD4 T cells compared with their blood-derived counterparts, while *IL11RA* and *LIFR* were expressed at similar levels in both compartments (Fig. 5C). This upregulation was recapitulated when PBMC-derived CD4 T cells were exposed to tumor CM (Fig. 5D), suggesting that tumor-derived factors, likely IL-6, drive the transition from CD4.CTL to a Tfh-like phenotype within the TME. To test this hypothesis, we examined the effect of tumor CM on gene expression in KLRG1⁺ CD4 T cells. Exposure to CM promoted a Tfh-like phenotype in peripheral blood-derived KLRG1⁺ CD4 T cells, which was reversed by IL-6 receptor blockade (tocilizumab) (Fig. 5E).

These data support a role for the TME, likely mediated by IL-6 signalling via gp130, in modulating CD4 T cell cytotoxicity. Consistent with this, blockade of the IL-6 receptor in KLRG1^+^ CD4 T cells resulted in increased overall cytotoxic capacity (Fig. 5F), pointing to potential therapeutic strategies to enhance anti-tumor immune responses.

## DISCUSSION

CD8 T cells have traditionally been regarded as the primary cytotoxic population of the adaptive immune system, based on their capacity for direct tumor cell killing. However, recent insights into cytotoxic CD4 T cells (CTLs) have reshaped this view, highlighting their direct role in anti-tumor immunity [14]. Nevertheless, their defining phenotype and the molecular mechanisms underlying their cytotoxic activity remain incompletely understood.

Here, we integrate single-cell, spatial transcriptomics approaches and subcellular analyses by cryo-expansion microscopy to patients-derived cancer samples to define cytotoxic CD4 T cells in human tumors. Notably, we found that CD4 CTL-mediated cytotoxicity does not rely exclusively on classical cytotoxic molecules but is instead associated with a distinct molecular signature comprising NKG7, GZMB, GNLY and, importantly, KLRG1 as a defining surface of this subset.

KLRG1 has been associated with enhanced cytotoxic function in CD4, CD8 and NK cells, including increased production of IFN-γ, TNF-α and granzymes compared with KLRG1^-^ T cells, in the context of primary biliary cholangitis, inclusion body myositis [61, 62], and more recently bladder cancer [63]. Both conventional cytotoxic CD8 T cells and highly differentiated cytotoxic CD4 T cells express KLRG1 following primary hCMV infection [64]. Moreover, the mRNA expression of KLRG1 in CD4 T cells across multiple human cancer types, including melanoma [14] and colorectal cancer [12], supports its role as a marker for cytotoxic CD4 T cells in tumor settings.

In contrast, KLRG1^+^ CD4 T cells have also been observed in murine colorectal cancer [65] and human renal cancer [66, 67], where they were associated with tumor progression. However, in those studies, KLRG1^+^ CD4 T cells were defined by a marker combination of Treg phenotypes and were not linked to a cytotoxic CD4 T cell program. In our study, by contrast, the Treg cluster, both in peripheral blood and tumor-infiltrated tissue, expressed KLRG1 at very low levels.

To further elucidate the cytotoxic mechanisms employed by CD4 CTLs, we used CRISPR/Cas9- mediated gene editing in primary human CD4 T cells to disrupt several candidate effector molecules. Among these, only deletion of GNLY, NKG7 and GZMB significantly impaired target cell killing. GNLY, a pore-forming protein, is initially produced as a 15-kDa precursor and subsequently processed into 9- kDa forms [68] with tumoricidal [69] and antimicrobial [70] properties within cytotoxic granules. Its presence in CD4 CTLs underscores a potential role in targeting bacterial infections not effectively controlled by classical cytotoxic cells [71, 72], and possibly tumor cells resistant to CD8 T cell or NK cell-mediated killing.

NKG7, an essential regulator of granule exocytosis [73], typically associated with NK cells, is expressed in CD4 TEMRA cells alongside GZMB and GNLY, indicating a robust cytotoxic machinery [34]. Together, these observations suggest that tumor-specific human CD4 CTLs may employ a coordinated mechanism in which GNLY mediates pore formation, NKG7 facilitates vesicle fusion and GZMB induces apoptosis in target cells. Future studies should investigate the temporal dynamics of this sequence in live cells, for example using high-throughput imaging approaches such as spinning disk microscopy. Such analysis may help explain the differences that we observed in killing kinetics and contact duration between CD4 CTLs and CD8 T cells in our single-cell picowell assays (Fig. 1A, right).

Our findings also provide insight into the interaction between cytotoxic CD4 T cells and tumor cells, mediated through ligation of N-cadherin (CD325) [58]. This interaction contributes to the formation and stabilization of the immunological synapse, as previously described for E-cadherin (CD324) [74], and may be essential for enabling KLRG1^+^ CD4 T cells to exert their cytotoxic functions, targeting and eliminating tumor cells. This mechanism is reminiscent of intercellular adhesion molecule 1 (ICAM-1), which enhances immunological synapse stability and is critical for effector and memory responses [75]. In line with this, activation of ICAM-1–deficient CD8 T cells *in vivo* results in an increased proportion of KLRG1^+^ T cells [76]. KLRG1 expression may therefore confer cytotoxic potential to CD4 T cells by influencing the stability of the interaction with their targets and/or that only CD4 T cells with TCRs in a defined range of affinity might naturally upregulate KLRG1 to acquire cytotoxic features.

While such cytotoxic activity is beneficial for tumor targeting, it would be detrimental if directed against APCs. Indeed, in murine models, Tr1 CD4 cells expressing IL-10, GZMB, PRF1, CCL5 and LILRB4, can impair anti-tumor immunity by killing cDC1s [59]. In contrast, our data in human systems demonstrate that KLRG1 ligands are absent on APCs, suggesting that collateral damage to non-tumor MHC class II^+^ cells is minimal, as we functionally confirmed *in vitro* using human B cells as APCs.

KLRG1 thus acts as a molecular checkpoint that enables selective targeting of cadherin-expressing tumor cells while sparing cadherin-negative APCs. This provides a mechanism of APC protection and addresses the long-standing question of how CD4⁺ T cells can exert cytotoxicity without compromising immune homeostasis.

In the context of cancer, melanoma progression is instead characterised by a “cadherin switch”, involving loss of E-cadherin, typical of normal melanocytes, and the acquisition of N-cadherin, which promotes tumor cell detachment and interaction with stromal components [77]. Importantly, melanoma cells appear to depend on continued cadherin expression for survival and migration [78], suggesting that they cannot downregulate N-cadherin to evade CD4 CTL-mediated killing without compromising tumor fitness. Indeed, cadherin-negative melanomas are rare, supporting the idea that cadherin expression is essential for tumor maintenance [79]. These observations are encouraging in view of potential therapeutic applications of our results, since MHC class II^+^ APCs within and outside the TME would be preserved and could present antigens, while tumor cells would be efficiently eliminated.

Despite these promising findings, the reduced abundance of the CD4 CTL cluster in TILN and their compromised cytotoxicity within the TME signal significant hurdles both for natural anti-tumor responses and for harnessing them therapeutically. This observation might suggest either a compromised capacity of CD4 CTLs to infiltrate the TME, or an in-situ conversion of the penetrated cells towards a Tfh-like phenotype.

Tfh differentiation is known to depend on IL-6 in combination with IL-21 [80]. In the absence of these cytokines, Tfh development is impaired [81], whereas either cytokine alone is insufficient to drive differentiation [82]. Consistent with this, the elevated levels of IL-6 secreted by tumor cells, together with the upregulation of the IL-6R, support an inhibitory effect of this cytokine on CD4 CTLs, once they enter the TME. In multiple contexts, inhibition of IL-6 signalling has emerged as a promising strategy to enhance anti-tumor immunity [83–85], raising the possibility that such approaches may also limit CD4 CTL plasticity towards Tfh states, as supported by our *in vitro* data.

From an ACT perspective, future work may focus on engineering KLRG1^+^ CD4 T cells to express CARs or high-affinity tumor-specific TCRs, thereby enhancing their tumor-targeting capacity. However, despite their promising application in ACT and potential advantageous role in eliminating tumors [29], caution is warranted, as in multiple myeloma, patients with limited therapeutic responses exhibited higher frequencies of cytotoxic CD4 CAR T cells [86].

Finally, although our study was restricted to treatment naïve melanoma patients, further work will be required to validate these signatures and cytotoxic mechanisms across additional solid and haematological tumors, including longitudinal analyses before and after treatment. Initial insights on the potential predictive/prognostic value of CD4 CTLs in the response to immune checkpoint blockade stem from the recent observation that a CD4 CTL intratumoral gene signature is predictive of anti-PD-L1 treatment efficacy in patients with metastatic bladder cancer [13].

## FIGURE LEGENDS

SUPP. FIGURE 1

(A) Characteristics of melanoma patients included in this study. (B) UMAP of the integrated total CD4^+^ T cell scRNA-seq dataset colored by unsupervised graph-based clusters. (C) Dot plot of canonical marker genes across the unsupervised clusters shown in (B); dot size indicates the fraction of expressing cells and color indicates per-gene scaled average expression. (D) Concordance between manual CD4 state annotation and reference-derived labels: stacked bar plot shows, for each manual state, the proportion of cells assigned to each ProjecTILs CD4 reference-map functional label. (E) Cytotoxic program strength across tissues: per-sample mean UCell cytotoxic signature scores for the Cytotoxic CD4^+^ population summarized by tissue (PBMC vs TILN). (F) Projection of tumor antigen– specific CD4^+^ T cells onto the integrated total CD4^+^ reference embedding; projected cells are overlaid on the reference atlas to visualize state correspondence. (G) Multi-cancer cohort: heatmap showing scaled expression of representative gene programs for cytotoxic and Tfh across samples, annotated by tissue of origin and cancer type. Cancer types: AML (Acute Myeloid Leukaemia), BM (Bone Marrow), BCC (Basal Cell Carcinoma), BCL (B-cell Lymphoma), BC (Breast Cancer), CRC (Colorectal Cancer), ESCA (Esophageal Cancer), FTC (Follicular Thyroid Carcinoma), HCC (Hepatocellular Carcinoma), LUNG (Lung Cancer), CHOL (Cholangiocarcinoma), Melanoma, MM (Multiple Myeloma), NPC (Nasopharyngeal Carcinoma), OV (Ovarian Cancer), PACA (Pancreatic Adenocarcinoma), RC (Rectal Cancer), SCC (Squamous Cell Carcinoma), THCA (Thyroid Cancer), UCEC (Uterine Corpus Endometrial Carcinoma). (H) Multi-cancer cohort: top 20 marker genes of the cytotoxic CD4 state versus all other CD4 T cells ranked by log2 fold change, with genes categorized by predicted cell-surface receptor status versus other genes.

SUPP. FIGURE 2

(A) Supernatant cytokine quantification (pg/ml) after killing, paired t-test (n=4). (B) Cryo-expansion microscopy of CD4 and CD8 T cells labelled for LAMP1 (green), NHS (blue) and GNLY (magenta), scale: 1 µm. Bottom panel: quantification of GNLY per cell, granule volume and diameter represented using Tukey box and whiskers plot with median (in-box line) and outliers (dots), Mann-Whitney unpaired t-test.

SUPP. FIGURE 3

(A) STRING protein–protein interaction network centered on KLRG1 in PBMC CD4 T cells. Only proteins directly connected to KLRG1 are displayed. Node size reflects the absolute log2 fold change (|log2FC|) from differential expression analysis (Fig. 2B), and edge width corresponds to the STRING combined confidence score. KLRG1 is highlighted in red, while adhesion-related interaction partners are shown in blue. (B) Representative example of the expression of CD324 and of CD325 by flow cytometry on melanoma cell lines and on tumor cells from TILN. (C) Gating strategy for the identification of cDC1, cDC2, pDC, B cells and monocytes.

SUPP. FIGURE 4

(A) Summary of regions of interest (ROIs) used for spatial transcriptomics analysis. The table reports, for each patient, the number of CD4⁺ and CD20⁺ ROIs sampled inside or outside the tumor region, as well as CD4⁺ ROIs classified by proximity to B cells (close or far). (B) Pseudobulk-normalized expression of *IFI6* and *MX1* across annotated CD4⁺ T cell subsets in PBMC and TILN samples from the total CD4^+^ scRNA-seq cohort. Expression values were aggregated per patient and tissue, with boxplots summarizing patient-level distributions and points representing individual samples.

