## Supplementary material for "KLRG1 identifies circulating cytotoxic CD4 T cells with selective anti-tumor function in human cancer": SUPP Figure 1

**A** Melanoma patients characteristics included in this study

| Patient | | | Diagnosis | | | Antigen expression | | |
|---|---|---|---|---|---|---|---|---|
| ID | Sex | HLA allele | TNM | Stage | Age | Mel-A | ESO-1 | Mage3 |
| LAU 706 | F | DRB3*02:02 | pTx N0 M0 | III | 64 | - | + | ++ |
| LAU 380 | F | DRB1*07:01 | pTx N1b M0 | III | 63 | +++ | - | - |
| LAU 149 | F | DRB1*07:01 | pT2b N0 M0 | IIA | 19 | ++ | + | ++ |
| LAU 42 | M | DPB1*04:01 | pT4a N0 M0 | III | 47 | + | + | +++ |
| LAU 1417 | M | DPB1*04:01 | pT3bp N2 M0 | IIIA | 66 | +++ | +++ | +++ |
| LAU 372 | M | DPB1*04:01 | T4a N2 M0 | III | 59 | - | - | - |
| LAU 1397 | F | DRB1*07:01 | Txc N3c M0 | IIIC | 22 | +++ | -/+ | +++ |
| LAU 50 | M | DPB1*04:01 | pT4a N0 M0 | IIB | 62 | +++ | + | ++ |
| LAU 969 | M | DPB1*04:01 | pT3b N2 M0 | IIIB | 50 | +++ | - | + |

**B** UMAP of the total CD4 T cell scRNAseq dataset

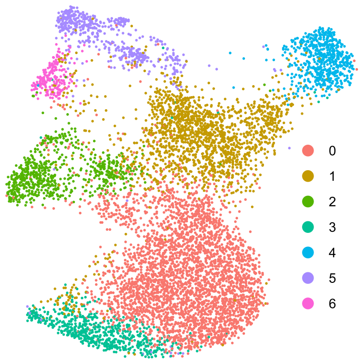


**C** Canonical marker genes in total CD4 T cells

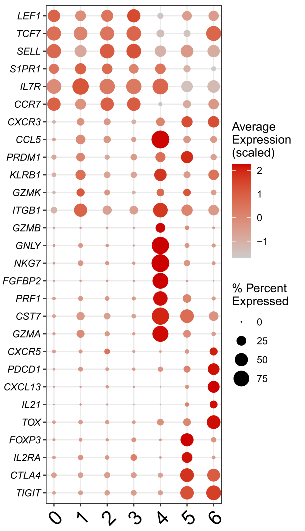


**D** Manual vs reference-derived CD4 state annotation

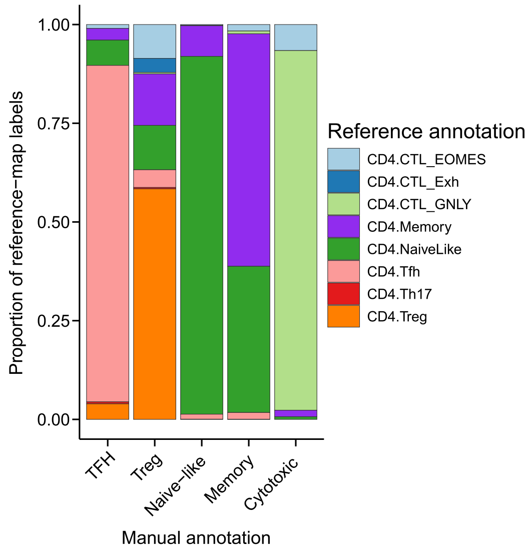


**E** Cytotoxic program across tissues

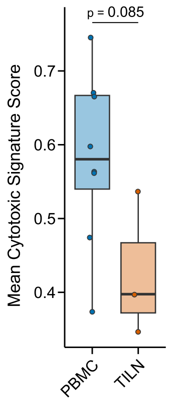


**F** Tumor-specific CD4 T cell projection

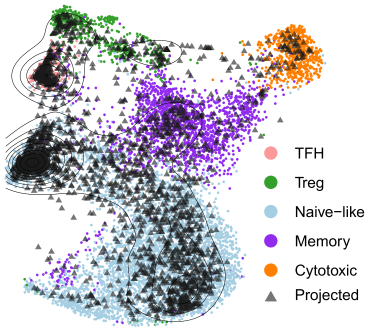


**G** Multi-cancer cohort

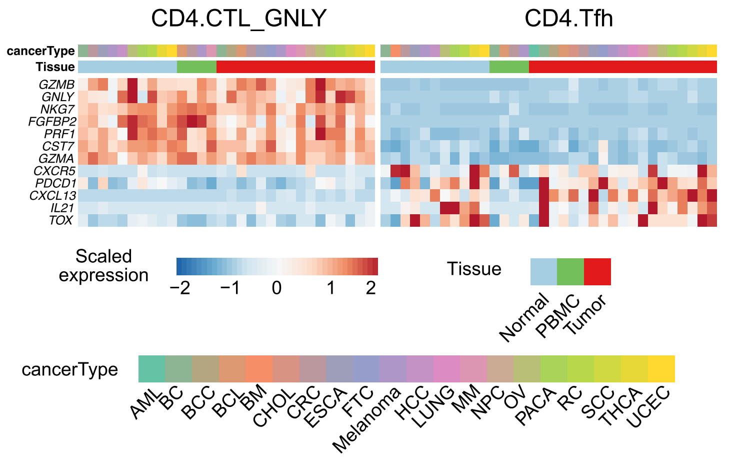


**H** Top cytotoxic genes in multi-cancer cohort

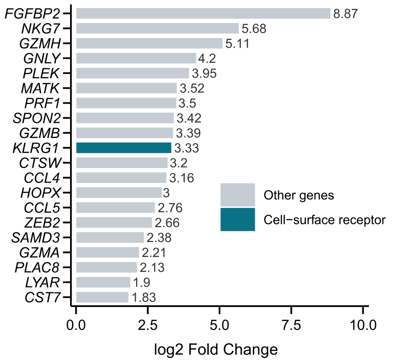
