## Supplementary material for "KLRG1 identifies circulating cytotoxic CD4 T cells with selective anti-tumor function in human cancer": SUPP Figure 2

A

Supernatant cytokine quantification after killing

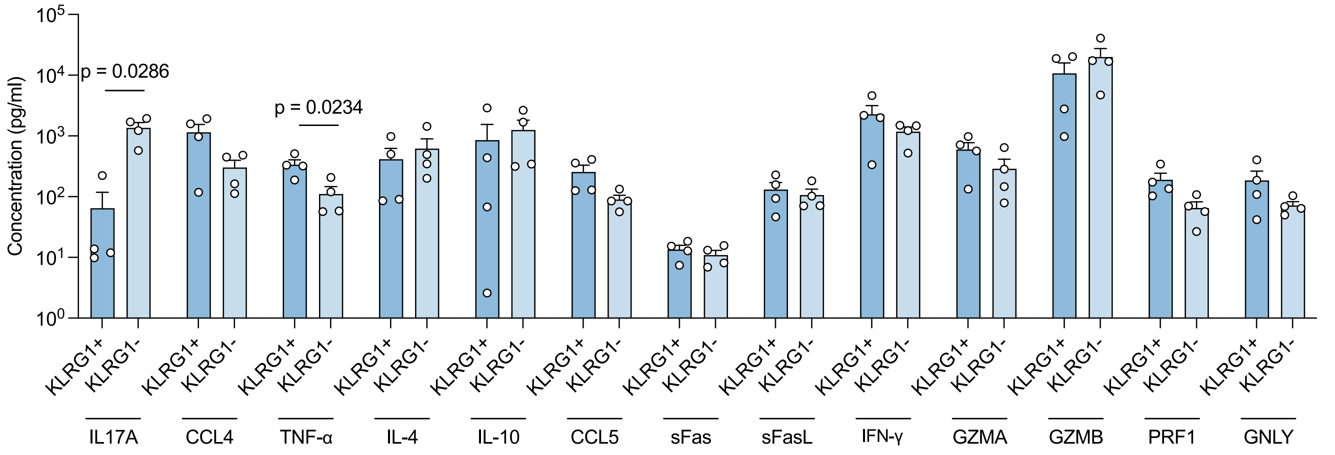


B

Cryo-expansion microsopy of GNLY in CD4 CTL vs CD8 T cell

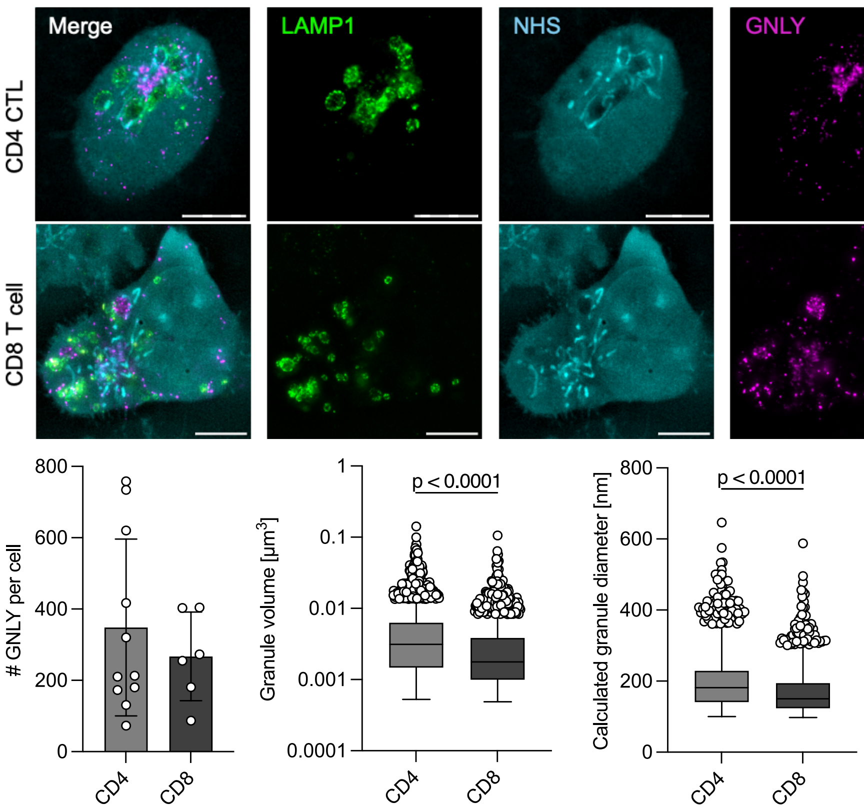
