## Supplementary material for "KLRG1 identifies circulating cytotoxic CD4 T cells with selective anti-tumor function in human cancer": SUPP Figure 3

A KLRG1-adhesion interaction network

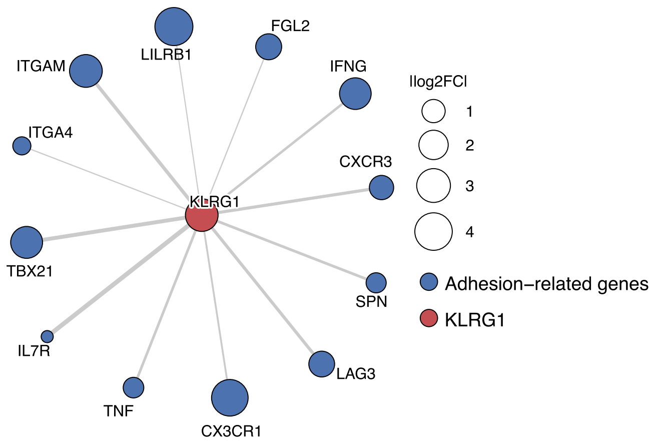


B CD324 and CD325 expression by flow cytometry

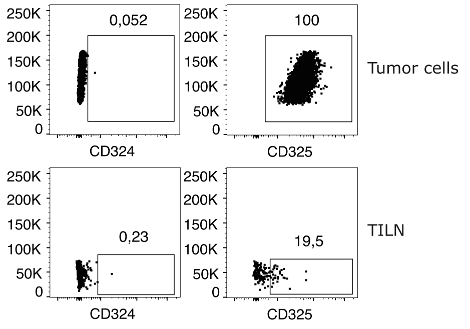


C Gating strategy for APCs

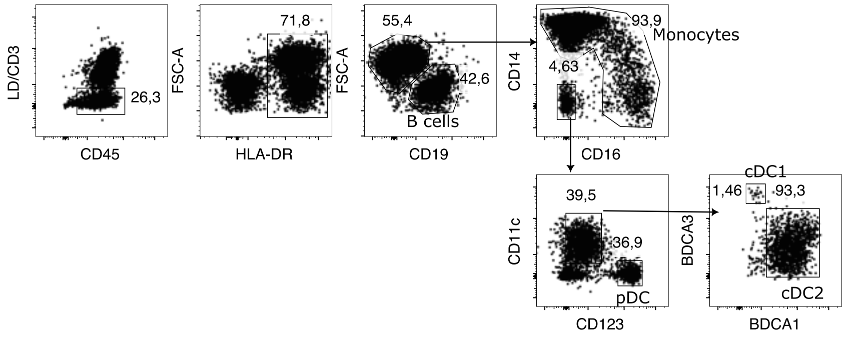
