## Supplementary material for "KLRG1 identifies circulating cytotoxic CD4 T cells with selective anti-tumor function in human cancer": SUPP Figure 4

A

Summary of regions of interest used for spatial transcriptomics analysis

| | CD4+ enriched | | | | CD20+ enriched | | |
|---|---|---|---|---|---|---|---|
| | Tumor region distance | | CD20+ spot distance | | Tumor region distance | | |
| Patient | Inside | Outside | Close | Far | Inside | Outside | Total spots |
| LAU706 | 8 | 2 | 3 | 7 | 2 | 0 | 12 |
| LAU380 | 4 | 2 | 6 | 0 | 4 | 2 | 12 |
| LAU1417 | 4 | 1 | 5 | 0 | 0 | 7 | 12 |
| LAU372 | 8 | 4 | 0 | 12 | 0 | 0 | 12 |
| LAU1397 | 5 | 1 | 6 | 0 | 5 | 1 | 12 |

B

Gene expression in total CD4 in scRNAseq cohort

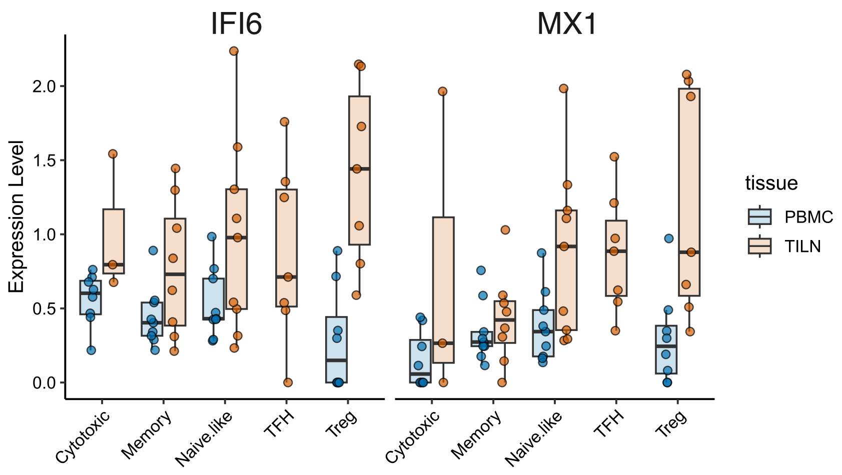
